# Nonlinearities between inhibition and T-type calcium channel activity bidirectionally regulate thalamic network oscillations

**DOI:** 10.1101/2020.06.02.129601

**Authors:** Adam C Lu, Christine K Lee, Max Kleiman-Weiner, Brian Truong, Megan Wang, John R Huguenard, Mark P Beenhakker

**Affiliations:** Department of Pharmacology, University of Virginia, Charlottesville, VA, USA; Department of Neurosurgery, Massachusetts General Hospital, Boston, MA, USA; Department of Psychology, Harvard University, Cambridge, MA, USA; Princeton Neuroscience Institute, Princeton University, Princeton, NJ, USA; Department of Neurology, Stanford University, Palo Alto, CA, USA

## Abstract

Absence seizures result from 3-5 Hz generalized thalamocortical oscillations that depend on highly regulated inhibitory neurotransmission in the thalamus. Efficient reuptake of the inhibitory neurotransmitter GABA is essential, and reuptake failure worsens seizures. Here, we show that blocking GABA transporters (GATs) in acute brain slices containing key parts of the thalamocortical seizure network modulates epileptiform activity. As expected, we found that blocking either GAT1 or GAT3 prolonged oscillations. However, blocking both GATs unexpectedly suppressed oscillations. Integrating experimental observations into single-neuron and network-level computational models shows how a non-linear dependence of T-type calcium channel opening on GABA_B_ receptor activity regulates network oscillations. Receptor activity that is either too brief or too protracted fails to sufficiently open T-type channels necessary for sustaining oscillations. Only within a narrow range does prolonging GABA_B_ receptor activity promote channel opening and intensify oscillations. These results have implications for therapeutics that modulate GABA transporters.

## Introduction

Neural circuits rely on a combination of intrinsic cellular properties and synaptic connections to generate large-scale electrical oscillations that drive behavior (Getting, 1989; Marder and Calabrese, 1996; Nusbaum and Beenhakker, 2002; Huguenard and McCormick, 2007). Following membrane hyperpolarization, such as that produced by synaptic inhibition, cortically-projecting neurons of the thalamus [i.e. thalamocortical (TC) neurons] produce brief bursts of action potentials (Llinás and Jahnsen, 1982), a cellular property that maintains both sleep-related and seizure-related oscillations (David A McCormick and Diego Contreras, 2001; Huguenard and McCormick, 2007; Beenhakker and Huguenard, 2009). Several studies have shown that CaV3.1 T-type calcium channels (T channels) sustain post-inhibitory rebound bursts in thalamocortical neurons by producing a relatively prolonged calcium-dependent, low-threshold spike (Kim et al., 2001, 2003; Porcello et al., 2003). These channels require membrane depolarization to open and hyperpolarization to recover (Coulter et al., 1989). Hyperpolarization-dependent recovery involves the removal of T channel inactivation (i.e. *de-inactivation*). As T channels are largely inactivated at resting membrane potentials, membrane hyperpolarization is necessary for robust rebound bursting (Llinás and Jahnsen, 1982; Coulter et al., 1989). While controlled voltage-clamp experiments have informed our understanding of how neuronal membrane potential dynamics can affect T channel opening (Gutierrez et al., 2001), we still know little regarding channel behavior during physiological forms of synaptic inhibition.

Reticular thalamic (RT) neurons serve as the main source of inhibitory, GABAergic input to thalamocortical neurons, especially in rodents (Shosaku, 1985; Pinault and Deschênes, 1998). Thalamocortical neurons express synaptic α_1_β_2_γ_2_ GABA_A_ receptors, and two types of extrasynaptic receptors: GABA_A_ (α_4_β_2_d) and GABA_B_ (Pirker et al., 2000; Kulik et al., 2002; Jia et al., 2005). Studies have shown that modulation of *synaptic* GABA_A_ receptors between RT and TC neurons has little effect on thalamocortical oscillations (Sohal et al., 2003; Rovó et al., 2014). In contrast, *extrasynaptic* receptors have been implicated in thalamocortical seizures, both for GABA_A_ (Cope et al., 2009) and GABA_B_ (Liu et al., 1992; Vergnes et al., 1997; Bortolato et al., 2010) receptors. Prior experimental and computational work has demonstrated that a shift from GABA_A_ receptor-mediated to GABA_B_ receptor-mediated inhibition at the RT-TC synapse transforms oscillations from a 10 Hz, sparse, spindle-like activity to a 3 Hz, hyper-synchronized, seizure-like state (von Krosigk et al., 1993; Destexhe et al., 1996; Destexhe, 1998; Blumenfeld and McCormick, 2000).

GABA transporters (GATs) powerfully control the activation of GABA_B_ receptors (Beenhakker and Huguenard, 2010). GAT1 and GAT3 represent the primary GABA transporters expressed in the brain and normally recycle GABA from the extrasynaptic space, thereby regulating GABA spillover from the synapse and the activation of extrasynaptic GABA_A_ and GABA_B_ receptors (Cope et al., 2009; Scanziani, 2000). In the thalamus, the more abundant transporter, GAT3, is localized farther away from synapses than GAT1 (De Biasi et al., 1998; Beenhakker and Huguenard, 2010). Consequently, specific GAT1 blockade results only in an increase in the amplitude of the GABA_B_ IPSC, reflecting increases in GABA concentration near the synapse. In contrast, specific GAT3 blockade results in an increase in both amplitude and decay of the GABA_B_-mediated inhibitory post-synaptic current (GABA_B_ IPSC), as GABA is allowed to diffuse far from the synapse where there is an abundance of GABA_B_ receptors (Beenhakker and Huguenard, 2010). On the other hand, dual GAT1 and GAT3 blockade results in a roughly 10-fold increase in the decay time constant and a 5-fold increase in the area under the curve of the GABA_B_ IPSC. These findings were replicated in a diffusion-based computational model (Beenhakker and Huguenard, 2010).

In this study, we investigate the consequences of physiologically-relevant GABA_B_ receptor-mediated inhibition observed during different combinations of GABA transporter blockade: control, GAT1 blockade, GAT3 blockade and dual GAT1+GAT3 blockade (Beenhakker and Huguenard, 2010). As absence seizures are dependent on GABA_B_ receptor signaling, we hypothesized that GAT blockade would regulate both thalamocortical neuron rebound bursting and network-level thalamic oscillations. We examine the effects of different GABA_B_ receptor activation waveforms on both absence-seizure-like thalamic oscillations and single thalamocortical neuron responses. We first use pharmacological manipulations to demonstrate that individual GAT1 or GAT3 blockade prolongs seizure-like oscillations, but that dual GAT1+GAT3 blockade surprisingly abolishes oscillations. Next, we apply physiological GABA_B_ IPSC waveforms corresponding to each pharmacological condition to single thalamocortical neurons with dynamic clamp and demonstrate that individual GAT1 or GAT3 blockade increases rebound burst probability, but that dual GAT1+GAT3 blockade suppresses it. We then build computational model neurons to explore how the differential GABA_B_ IPSC modulation affects TC responses and discover that differential T channel gating dynamics are responsible for those differences. Finally, we build computational model thalamic networks to determine how the differential effects of GAT blockade influence thalamic oscillations and to probe for pro- and anti-epileptic mechanisms. Through these experimental and computational approaches, we identify how GABA_B_-mediated inhibition across both voltage and time dimensions regulates T channel activity and seizure-like oscillations.

## Results

### Thalamic oscillations

To evaluate the contribution of GABA transporters to thalamic network activity in the context of GABA_B_ receptor-mediated inhibition, we used a standard, acute rat brain slice model in which electrical oscillations are evoked by extracellular stimulation of the synaptic inputs to the reticular thalamic nucleus in the presence of the GABA_A_ receptor blocker bicuculline (Huguenard and Prince, 1994; Jacobsen et al., 2001; Kleiman-Weiner et al., 2009). We evoked oscillations at intervals producing no rundown [once per minute, (Jacobsen et al., 2001)] and monitored neuronal activity with extracellular multiunit electrodes placed within the ventrobasal complex of the thalamus. By detecting evoked bursts, we found that oscillations last between 2-13 seconds at baseline (Figure 1A). The autocorrelogram of binned spike times revealed pronounced secondary peaks at multiples of approximately 500 ms (Figure 1A). After recording evoked oscillations for 20 minutes under baseline conditions, we then applied one of four experimental solutions to the perfusate (Figure 1B). Experimental solutions consisted of: (1) a control solution identical to the baseline solution, (2) 4 µM NO-711, a specific GAT1 blocker (Sitte et al., 2002), (3) 100 µM SNAP-5114, a specific GAT3 blocker (Borden et al., 1994), or (4) a combination of 4 µM NO-711 and 100 µM SNAP-5114. These blocker concentrations achieve full GAT blockade (Beenhakker and Huguenard, 2010). Experimental solutions were applied for 40 minutes, and then washed out over another 20 minutes.

**Figure 1.**
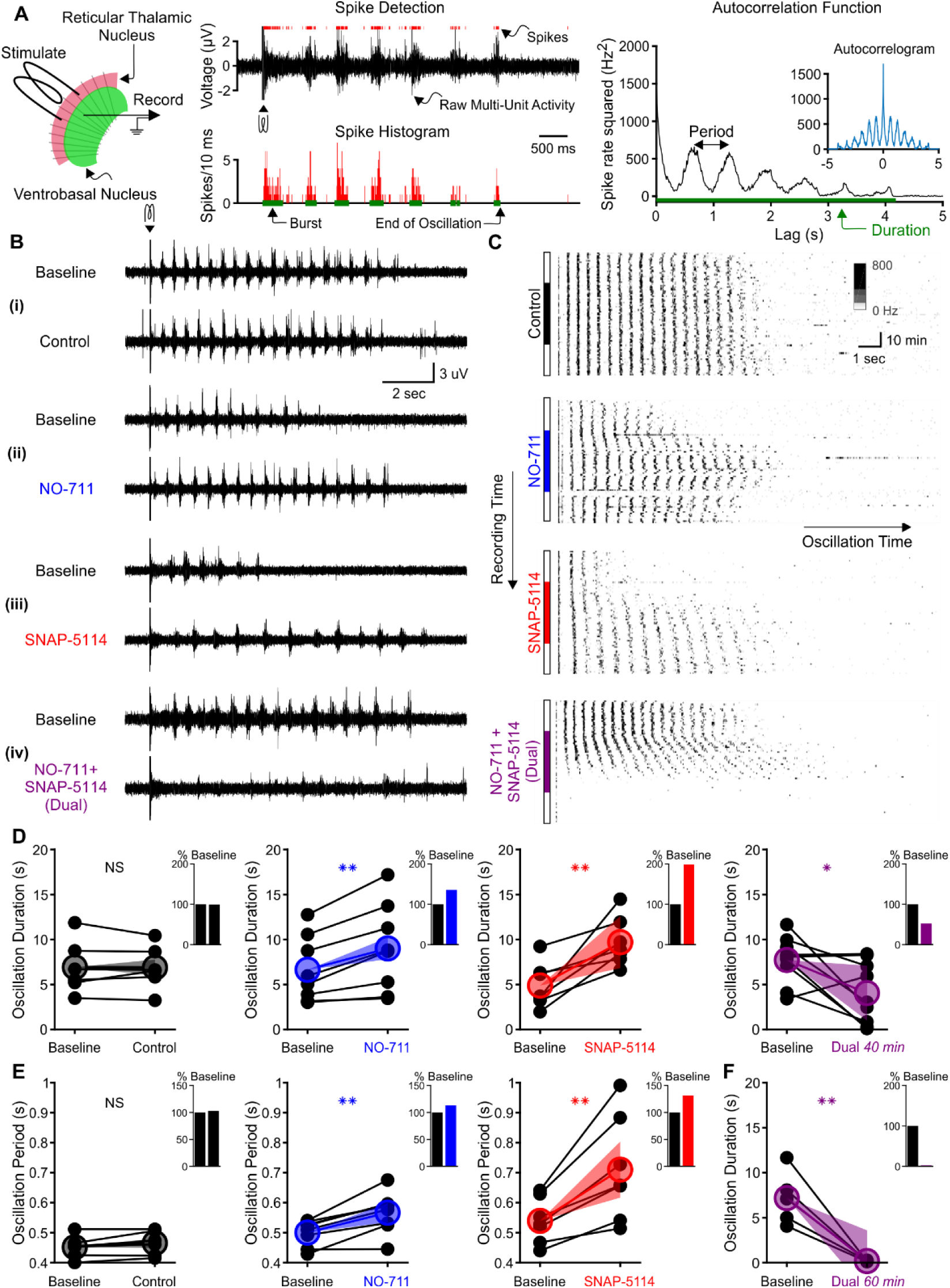
Individual GAT1 or GAT3 blockade strengthens thalamic oscillations, but dual GAT1+GAT3 blockade abolishes oscillations. (**A**) Slice recording setup and sample analysis. Acute thalamic slices were bathed in bicuculline to block GABA_A_ receptors. A brief voltage stimulus (0.5 ms, 10 V) was applied with a bipolar electrode placed in either the reticular thalamic nucleus or the adjacent internal capsule to evoke epileptiform oscillations recorded extracellularly in the ventrobasal complex. Spikes were detected, binned, and grouped into bursts. The oscillation duration was measured from the spike histogram and the oscillation period was computed from the autocorrelation function of binned spikes. (**B**) Example evoked epileptiform oscillations at baseline and 40 minutes after perfusing with (i) control (no drug added), (ii) 4 µM NO-711 (GAT1 blocker), (iii) 100 µM SNAP-5114 (GAT3 blocker) or (iv) dual 4 µM NO-711+100 µM SNAP-5114. (**C**) Example PSTHs over the entire course of a single control, GAT1-, GAT3-, and dual-block experiment. Oscillations were evoked every 60 seconds, but only the first 17 seconds after stimulation are shown. Drugs were perfused for 40 minutes after 20 minutes of baseline, followed by 20 minutes of washout. (**D-F**) Oscillation measures for all slices. Colored circles denote the mean value, colored lines denote the mean change; and shaded areas denote the 95% confidence intervals for the mean change. (**D**) Oscillation duration did not change following control perfusion, but increased following NO-711 or SNAP-5114 perfusion. Dual NO-711+SNAP-5114 perfusion reduced oscillation duration (* p < 0.05, ** p < 0.01, paired-sample *t*-test). Inset shows mean change relative to baseline. (**E**) Oscillation period did not change following control perfusion and lengthened following NO-711 or SNAP-5114 perfusion (** p < 0.01, paired-sample *t*-test). (**F**) After 60 minutes of dual NO-711 + SNAP-5114 perfusion, oscillations were abolished in all slices (** p < 0.01, paired-sample *t*-test).

When individually applied, either GAT1 or GAT3 blockade prolonged oscillations (Figure 1C), consistent with the absence seizure exacerbation seen with a clinically used GAT1 blocker, tiagabine (Ettinger et al., 1999; Knake et al., 1999; Vinton et al., 2005). We measured the duration of each evoked oscillation, then computed the average duration of the last 5 stable oscillations in baseline and experimental solutions (Figure 1D). Relative to baseline, individual GAT1 or GAT3 blockade increased oscillation duration by 36% (n = 8 slices from 5 animals, p = 0.0027) and 99%, (n = 7 slices from 5 animals, p = 0.0076), respectively. We also evaluated the effects of individual GAT1 or GAT3 blockade on the period of evoked oscillations (Figure 1E). Relative to baseline, GAT1 or GAT3 blockade increased the oscillation period by 13% (n = 8 slices from 5 animals, p = 0.0021) and 32% (n = 7 slices from 5 animals, p = 0.0050), respectively. Collectively, the effects of GAT blockade on oscillation properties generally agreed with the previously reported actions of GAT blockers on isolated GABA_B_ receptor-mediated IPSCs. That is, the 1.4-fold and 2-fold increase in oscillation duration corresponds roughly to the reported 1.5-fold and 2.2-fold increase in GABA_B_ IPSC amplitude produced by GAT1 or GAT3 blockade, respectively (Beenhakker and Huguenard, 2010). Additionally, GAT3 blockade significantly prolonged oscillation period, while the effect for GAT1 blockade was modest, consistent with reported effects on isolated GABA_B_ IPSC decay (Beenhakker and Huguenard, 2010).

Surprisingly, the effects of NO-711 and SNAP-5114 co-perfusion on evoked oscillations were not additive. Rather than prolonging evoked oscillations, dual GAT1+GAT3 blockade ultimately *eliminated* oscillations. Following a brief prolongation during the early phases of drug perfusion (see Figure 1C), dual GAT1+GAT3 blockade eventually decreased oscillation duration by 48% (Figure 1D, n = 9 slices from 4 animals, p = 0.026; here and in all results, percentages refer to relative change from control conditions). As the effects of dual blockade on oscillation duration did not appear to reach a steady state by 40 minutes, we extended the drug application to 60 minutes in a subset of experiments. For those slices, dual GAT1+GAT3 blockade invariably abolished oscillations (Figure 1F, n = 5 slices from 3 animals, p = 0.0062).

In summary, the observed effects of individual GAT1 or GAT3 blockade on oscillation duration and period generally reflect the actions the individual blockers have on GABA_B_ receptor-mediated IPSCs isolated from thalamocortical neurons. GAT blockade-dependent increases in IPSC amplitude were associated with increased strength of oscillation, as measured by duration. However, the effects of dual GAT1+GAT3 blockade on oscillation duration did not reflect the additive effects of combined blockade on GABA_B_ IPSCs (Beenhakker and Huguenard, 2010). To better understand the discrepancy between GAT regulation of IPSCs and GAT regulation of thalamic oscillations, we next examined how IPSC amplitude and kinetics regulate the activity of single thalamocortical neurons.

### Single neuron recordings

We investigated the effects of GAT-modulated, GABA_B_ receptor-mediated currents on thalamocortical neuron rebound bursting, as this property is likely critical for the initiation of each successive cycle of an evoked oscillation (von Krosigk et al., 1993; Huguenard and Prince, 1994; Warren et al., 1994). Experimentally evoked GABA_B_ receptor-mediated IPSCs isolated in acute thalamic slices vary considerably in amplitude (Beenhakker and Huguenard, 2010), likely reflecting differences in synaptic activation of reticular thalamic neurons by the electrical stimulus. We therefore utilized an alternative approach to systematically examine the effects of GAT blockade on the firing properties of thalamocortical neurons: dynamic clamp (Sharp et al., 1993; Ulrich and Huguenard, 1996). We used GABA_B_ receptor-mediated IPSC waveforms isolated under voltage clamp during each pharmacological condition (control, GAT1 blockade, GAT3 blockade, dual GAT1+GAT3 blockade) as conductance waveform commands applied to single thalamocortical neurons (Figure 2A). We refer to these dynamic clamp-mediated conductance waveforms as *d*IPSCs. We applied each *d*IPSC pharmacological condition (*d*Control, *d*GAT1-Block, *d*GAT3-Block, *d*Dual-Block; Figure 2B) to each recorded thalamocortical neuron.

**Figure 2.**
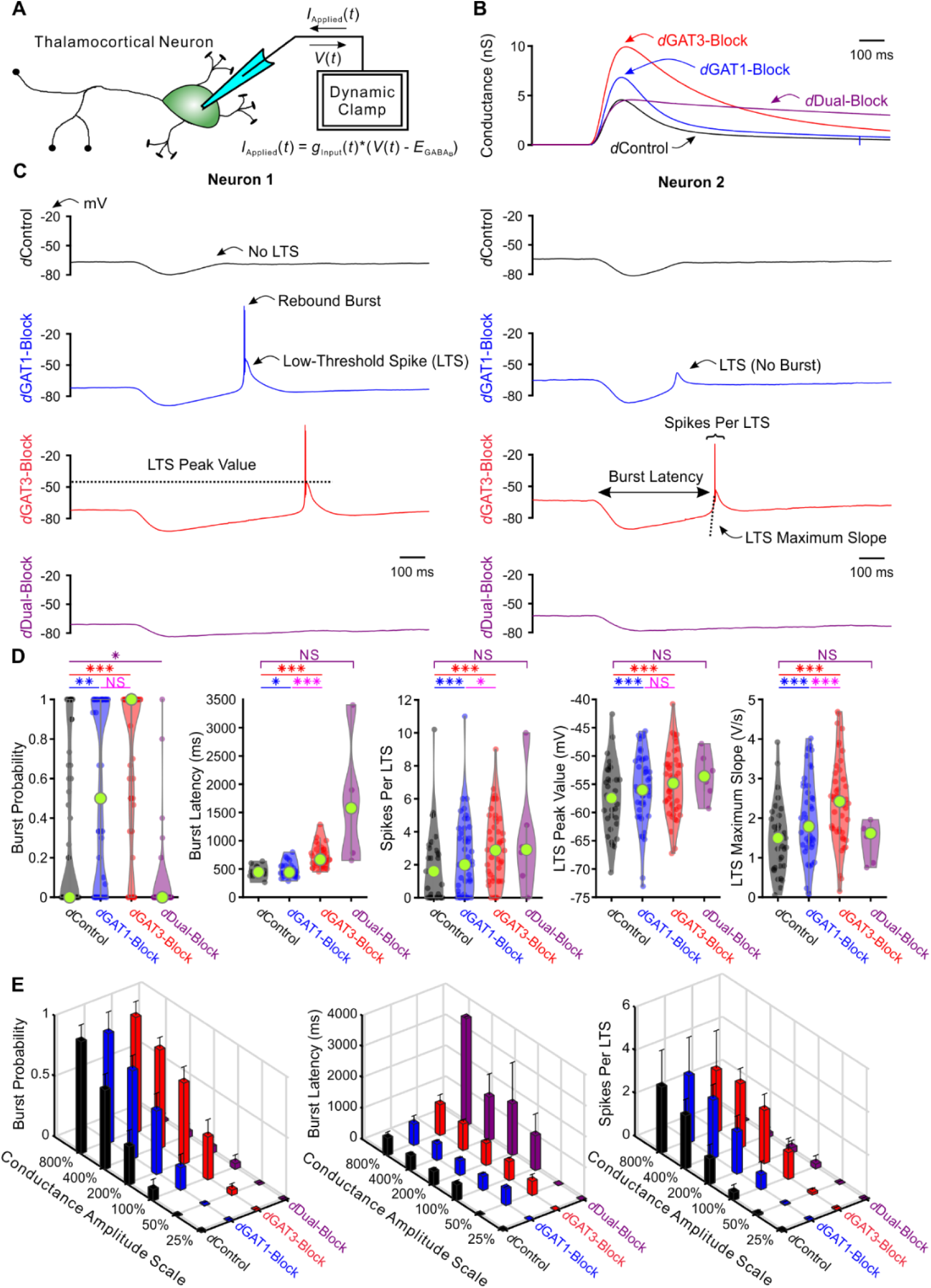
Post-inhibitory, low-threshold rebound spikes and bursts in thalamocortical neurons are bidirectionally modulated by GABA_B_ receptor-mediated conductance waveforms. (**A**) Dynamic clamp setup. A thalamocortical neuron was patched in the whole-cell configuration. The applied current was computed from the instantaneous voltage and a command conductance waveform over time to simulate GABA_B_ receptor activation. (**B**) Command GABA_B_ receptor conductance waveforms (*d*IPSCs, amplitudes scaled by 200%) that emulated different GAT blockade conditions based on GABA_B_ IPSCs isolated with voltage clamp (Beenhakker and Huguenard, 2010). (**C**) Sample voltage responses of two neurons to the four different *d*IPSCs shown in (B). Annotations are for measures in (D) and (E). (**D**) Distributions of post-inhibitory low-threshold rebound spike or burst measures over all 47 recorded neurons across *d*IPSCs shown in (B). Relative to *d*Control responses, rebound burst probability increased following either *d*GAT1-or *d*GAT3-Block, but decreased following *d*Dual-Block (* p < 0.05, ** p < 0.01, *** p < 0.001, Friedman’s test for burst probability, repeated-measures ANOVA otherwise). (**E**) Mean LTS or burst measures over all 47 recorded neurons, across 4 different *d*IPSC waveforms and 6 different conductance amplitude scales. Error bars denote 95% confidence intervals.

Since neurons likely receive variable numbers of inhibitory inputs, we scaled the conductance waveform amplitudes for each pharmacological *d*IPSC by 25%, 50%, 100%, 200%, 400% and 800%, yielding 24 possible *d*IPSC waveforms (i.e. 4 pharmacological conditions x 6 amplitude scales). Additionally, we delivered each of the 24 waveforms at three approximate holding potentials: -60 mV, -65 mV or -70 mV. Five non-consecutive repetitions were performed for each *d*IPSC waveform and holding potential condition.

Figure 2C shows example responses to *d*IPSCs scaled by 200% delivered with dynamic clamp. Post-inhibitory, low-threshold calcium spikes (LTS) often followed each *d*IPSC, with sodium-dependent action potentials often crowning each LTS. Herein, *LTS* refers to the slow, broad (∼50 ms) event following inhibition. *Burst*, on the other hand, specifically refers to the collection of action potentials crowning the LTS. We quantified several properties of each post-inhibitory LTS and burst in response to each *d*IPSC, including the probability of occurrence and the latency from *d*IPSC onset. We also computed LTS features such as peak voltage value, maximum rising slope and the number of spikes per LTS, averaged across trials for each neuron.

We first examined LTS and burst probability distributions over all recorded neurons following delivery of *d*IPSCs (LTS: not shown, burst: Figure 2D, 2E). Considering only those *d*IPSCs scaled by 200% (Figure 2D), relative to *d*Control responses, LTS and burst probabilities were higher following either *d*GAT1-Block (n = 47 cells, LTS: +26%, p = 0.0080, burst: +63%, p = 0.0018) or *d*GAT3-Block (n = 47 cells, LTS: +39%, p = 0.0015, burst: +106%, p = 1.7 × 10^−7^), but lower following *d*Dual-Block (n = 47 cells, LTS: -88%, p = 4.8 × 10^−6^, burst: -82%, p = 0.030). We observed the same trend across pharmacological conditions for all other conductance scales (LTS: not shown, burst: Figure 2E). Not surprisingly, increasing the conductance scale produced an increase in LTS and burst probability for either the *d*Control, *d*GAT1-Block or the *d*GAT3-Block condition. However, both probabilities were consistently very low, below 6%, across all conductance scales following *d*Dual-Block IPSCs. These changes in thalamocortical neuron rebound burst probability parallel the prolonged oscillation duration observed following individual GAT1 or GAT3 blockade, and the decrease in oscillation duration following dual GAT1+GAT3 blockade (Figure 1D).

Next, we examined distributions of average LTS and burst latencies over neurons responsive to *d*IPSCs (LTS: not shown, burst: Figure 2D, 2E). We restricted this analysis to *d*GAT1- and *d*GAT3-Block IPSCs because the *d*Dual-Block IPSC did not reliably evoke LTSs. Considering *d*IPSCs scaled by 200% (Figure 2D), relative to *d*Control responses, average LTS latency was not significantly different following *d*GAT1-Block (n = 32 cells, p = 0.97), while average burst latency was modestly prolonged (+4.6%, n = 21 cells, p = 0.034). In contrast, *d*GAT3-Block IPSCs significantly prolonged both LTS (+53%, n = 32 cells, p = 3.7 × 10^−9^) and burst latency (+58%, n = 21 cells, p = 1.1 × 10^−9^). We observed the same trend across pharmacological conditions for all other conductance amplitude scales (Figure 2E). As the inter-burst interval (latency from last burst) separates each cycle of seizure-like oscillations, and is dominated by inhibition of TC cells (Bal et al., 1995), the above results are consistent with the increase in oscillation period upon following either individual GAT1 or GAT3 blockade, but a more pronounced effect for the latter (Figure 1E).

We also examined the distributions of effects on LTS features across neurons responsive to *d*IPSCs (Figure 2D). Relative to *d*Control responses, there was an increase in average number of spikes per LTS, average LTS peak value and average LTS maximum slope following either *d*GAT1-Block (n = 32, spikes per LTS: +62%, p = 1.3 × 10^−6^, peak value: 2.5 ± 0.5 mV, p = 1.4 × 10^−4^, maximum slope: +52%, p = 2.6 × 10^−9^) or *d*GAT3-Block (n = 32, spikes per LTS: +93%, p = 9.3 × 10^−7^, peak value: 3.3 ± 0.8 mV, p = 5.2 × 10^−4^, maximum slope: +82%, p = 1.4 × 10^−9^). As both greater LTS peak value and greater LTS maximum slope increase the likelihood for action potential generation, these changes were consistent with both the observed increase in spikes per LTS following either *d*GAT1-Block or *d*GAT3-Block and, secondarily, the prolongation of oscillation duration following either GAT1 or GAT3 blockade (Figure 1D). In contrast, dual GAT1+GAT3 blockade reduced burst probability and abolished oscillations (Figure 1F).

In summary, the bidirectional differences in rebound burst probability, burst latency and LTS features of single thalamocortical neurons in response to different GABA_B_ activation waveforms were in agreement with the bidirectional differences in thalamic oscillation duration and period following the corresponding pharmacological manipulations. That is, by ultimately regulating thalamocortical neuron bursting, GATs appear to powerfully control thalamic network oscillations through differential activation of GABA_B_ receptors.

### Single neuron models

We next sought to determine the essential components of the thalamocortical neuron that contributes to the differential *d*IPSC responses observed during our dynamic clamp experiments. We also sought to better understand the underlying channel dynamics contributing to the differential responses. Towards these ends, we established a conductance-based, multi-compartment, single neuron model for each of the 36 experimentally recorded thalamocortical neurons for which we had stable responses across all acquired conductance amplitude scales (Figure 3A).

**Figure 3.**
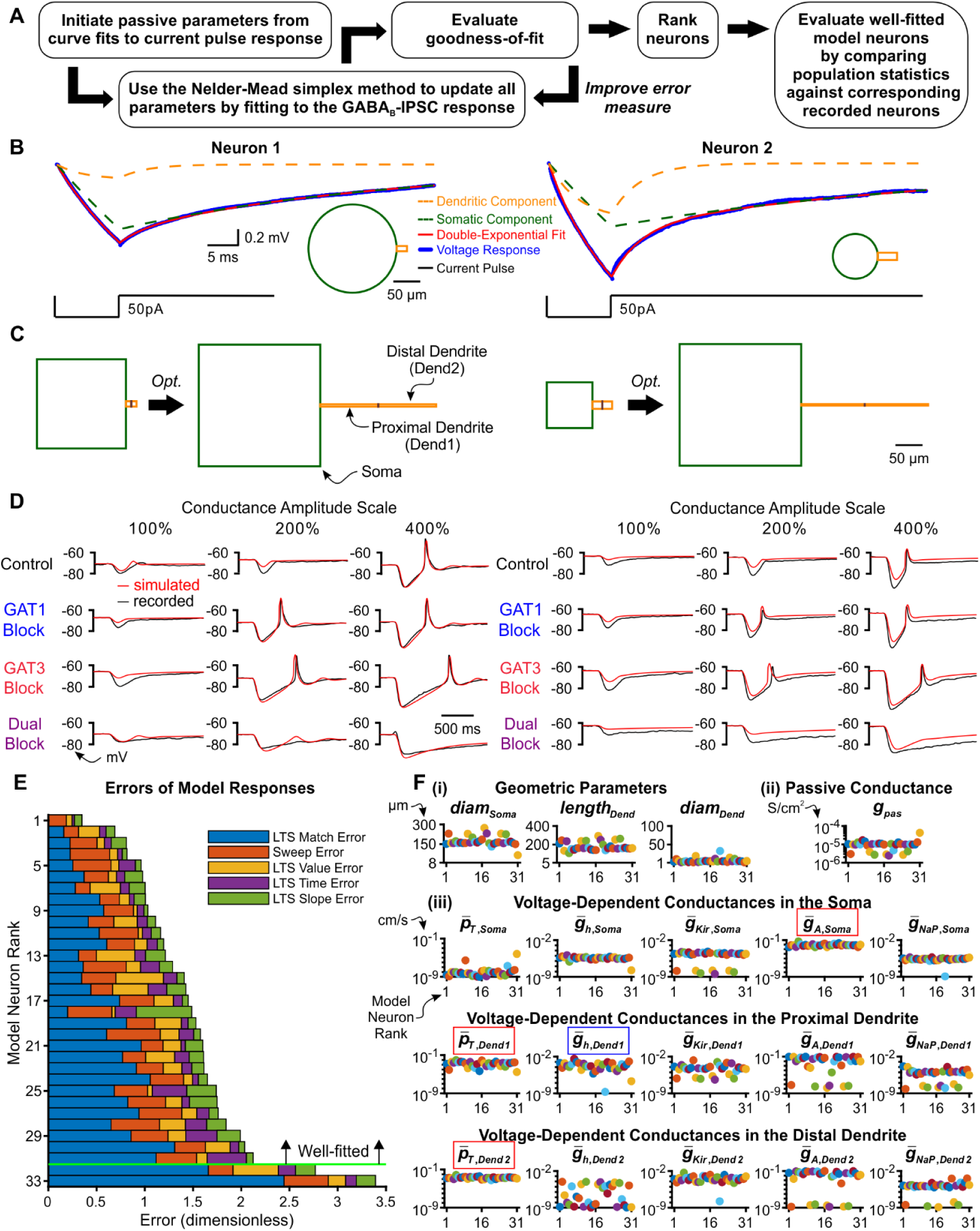
Thalamocortical model neurons reproduce GABA_B_ IPSC and rebound responses. (**A**) Model optimization workflow. (**B**) Sample double-exponential curve fits (red) to averaged current pulse responses (blue) for two example neurons. The dotted lines correspond to the curves representing the somatic compartment (green) and dendritic compartment (orange). The resulting ball-and-stick geometries estimated from the 2 exponential components are shown below the recordings (Johnston and Wu, 1994). (**C**) We converted ball-and-stick models shown in (B) to cylindrical, three compartment models (left), which were then optimized (right). (**D**) Fits of simulated *s*IPSC responses to recorded *d*IPSC responses, for the same two neurons. (**E**) The 33 model neurons that underwent optimization were ranked by a weighted average of 5 different types of errors (see *Methods*). The 31 highest ranked model neurons were considered *well-fitted*. (**F**) Final values of parameters that could vary for the 31 well-fitted model neurons. Note that the T channel density is high in the dendrites and the A-type potassium channel density is high in the soma for all model neurons (red boxes). The h channel density in the proximal dendrite negatively correlates with LTS latency (blue box). Geometric parameters are in µm, maximal conductance densities 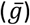 and conductance densities (*g*) are in S/cm^2^ and maximal permeability densities 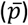 are in cm/s. The x axis is the model neuron rank in (E). We distinguish among (**i**) geometric parameters, (**ii**) voltage-independent *passive* parameters and (**iii**) voltage-dependent *active* parameters.

Our preliminary modeling results using existing TC cell models (Destexhe et al., 1998; Amarillo et al., 2014) failed to recapitulate two key features of GABA_B_ receptor-mediated post-inhibitory rebound LTSs that are likely critical in determining network level responses. Notably, the average LTS latencies of the model responses were routinely much earlier (400∼1000 ms) than the biological ones (400∼4000 ms). In addition, the model LTS and burst responses tended to be continuously graded in amplitude as a function of inhibitory strength, in contrast to the more characteristic all-or-none responses of recorded neurons. Therefore, we developed a gradient descent fitting approach to obtain suitable multicompartment models compatible with the data. As prior computational and experimental work demonstrates the importance of higher T channel densities in dendritic versus somatic compartments (Destexhe et al., 1998; Munsch et al., 1997; Williams and Stuart, 2000; Zhou et al., 1997), we modeled each thalamocortical neuron by a cylindrical somatic compartment and two cylindrical dendritic compartments in series (Figure 3C).

To reduce the number of fitted parameters, some simplifying assumptions were made: the somatic length and diameter were equivalent, the two dendritic compartments had equal dimensions, and passive leak channels were inserted into all three compartments at uniform densities. Four voltage-independent (passive) parameters were allowed to vary across model neurons: the somatic diameter (*diam*_*soma*_), the dendritic diameter (*diam*_*dend*_), the dendritic length (*length*_*dend*_) and the passive leak conductance density (*g*_*pas*_). As prior work has identified ionic currents that contribute to the resting membrane potential of thalamocortical neurons (Amarillo et al., 2014), we inserted the following five voltage-dependent channels in all three compartments: the T-type calcium channel (*I*_*T*_), the hyperpolarization-activated nonspecific cationic channel (*I*_*h*_), the A-type transient potassium channel (*I*_*A*_), the inward-rectifying potassium channel (*I*_*Kir*_) and the persistent sodium channel (*I*_*NaP*_). The densities of voltage-dependent channels were allowed to vary across compartments, resulting in a total of 15 (5 currents x 3 compartments) voltage-dependent (active) parameters that were allowed to vary across model neurons.

To provide an initial estimate of the geometric parameters that corresponded to each recorded thalamocortical neuron, we applied the short pulse methodology described by Johnston and Wu (Johnston and Wu, 1994, Chapter 4). During dynamic clamp experiments, we applied a short current pulse at the beginning of each recorded sweep. The average current pulse response for each neuron was well-fitted by a double exponential function (Figure 3B). From the coefficients and time constants of the two exponential components, we inferred the following four parameters for a ball-and-stick model (Rall, 1962): input conductance, electrotonic length, dendritic-to-somatic conductance ratio and the membrane time constant. These Rall model values were then converted into initial passive parameter seed values (*diam*_*soma*_, *diam*_*Dend*_, *length*_*dend*_, *g*_*pas*_) of each 3-compartment model neuron (see *Methods*).

Single thalamocortical neuron responses recorded during dynamic clamp experiments served to optimize passive and active parameters of each 3-compartment, thalamocortical model neuron. GABA_B_ receptors were placed in the somatic compartment of each model neuron, and activation waveforms identical to the conductance waveforms used in dynamic clamp (the *d*IPSCs) were applied. We refer to these simulated GABA_B_ receptor activation waveforms as *s*IPSCs, corresponding to each pharmacological condition (*s*Control, *s*GAT1-Block, *s*GAT3-Block, *s*Dual-Block). For each model neuron, simulated responses to *s*IPSCs were iteratively compared to experimental *d*IPSC responses. We evaluated the goodness-of-fit for each iteration with a total error defined by a weighted combination of component errors (see *Methods*). Examples of resultant geometry and voltage response fits are shown in Figures 3C and 3D, respectively.

Each model neuron was trained using a set of 12 recorded traces, each selected from a different *d*IPSC waveform, but evaluated against all recorded traces for the neuron and ranked by the total error (Figure 3E). By removing neurons with a total error greater than two standard deviations above the mean, we designated the top 31 neurons as the set of *well-fitted model neurons*. All well-fitted neurons were characterized by high T channel densities in the distal dendrite and high A-type potassium channel densities in the soma (Figure 3F). Considerable variability among model neurons was observed in the densities of other ionic channels, likely contributing to the heterogeneity in LTS and burst statistics among recorded neurons in response to each GABA_B_ *d*IPSC waveform (Figure 2D). For example, the value of the maximal h channel conductance density in the proximal dendrite 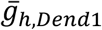 was negatively correlated with LTS latency (*R*^*2*^ = -0.80, not shown), which is consistent with the depolarizing effects of the h current (McCormick and Pape, 1990).

We compared the distribution of LTS probabilities and features over the 31 well-fitted model neurons and over their 31 corresponding neurons recorded in dynamic clamp. In general, there was high agreement between the model simulations and dynamic clamp recordings. We first compared the *d* and *s*IPSC datasets when scaled by 200% (Figure 4A and 4B). Relative to *d*/*s*Control responses, LTS probability was increased following *d/s*GAT1-Block (n = 31, model: +30%, p = 0.0086, recorded: +31%, p = 0.038) or *d/s*GAT3-Block (model: +44%, p = 7.4 × 10^−4^, recorded: +45%, p = 0.011) and decreased following *d/s*Dual-Block (model: -75%, p = 9.8 × 10^−7^, recorded: -93%, p = 5.7 × 10^−4^). Relative to *d/s*Control responses, average LTS latency was not different following *d/s*GAT1-Block (model: n = 27, p = 0.85, recorded: n = 21, p = 0.99) but was increased following *d/s*GAT3-Block (model: +42%, n = 27, p = 7.3 × 10^−8^, recorded: +52%, n = 21, p = 2.4 × 10^−6^). Differences in average number of spikes per LTS, average LTS peak value and average LTS maximum slope across *d*IPSC waveforms were not sufficiently captured by model neurons. The same trends across pharmacological conditions were observed for all other conductance amplitude scales, showing high agreement between model and recorded neurons for LTS probability and latency, but not for other LTS features (Figure 4C and 4D).

**Figure 4.**
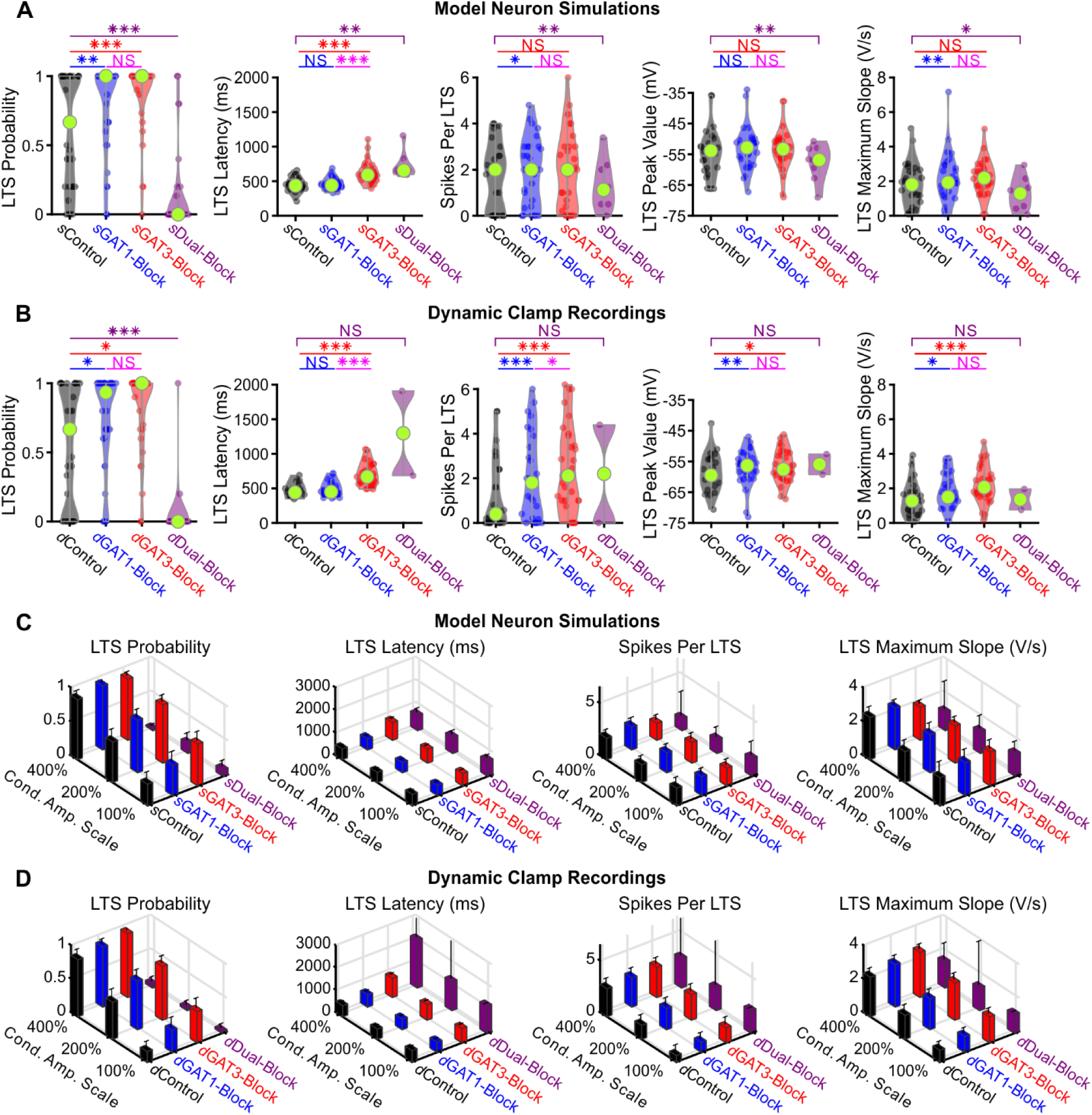
Well-fitted model and recorded neurons show similar low-threshold rebound spike differences in response to different GABA_B_ IPSC waveforms. (**A**) Distributions of post-inhibitory, low-threshold rebound spike measures over the 31 well-fitted model neurons across GABA_B_ IPSC waveforms shown in Figure 2B (* p < 0.05, ** p < 0.01, *** p < 0.001, repeated-measures ANOVA for LTS probability, Friedman’s test otherwise). (**B**) Same as (A) but for the corresponding 31 recorded neurons (Repeated-measures ANOVA for Spikes per LTS and LTS peak value, Friedman’s test otherwise). (**C**) Mean LTS or burst measures over all 31 model neurons, across 4 different GABA_B_ IPSC waveforms and 3 different conductance amplitude scales. Error bars denote 95% confidence intervals. (**D**) Same as (C) but for the corresponding 31 recorded neurons.

In summary, many single neuron models were established that sufficiently recapitulated the probability and timing of post-inhibitory rebound bursts in response to 12 different physiological GABA_B_-receptor IPSC waveforms. A commonality among well-fitted model neurons is that T channel densities were high in the dendrites and A-type potassium channel densities were high in the soma, while there was heterogeneity in other channel densities.

### Interplay between GABA_B_ receptors and T-type calcium channels

We next sought to understand the post-synaptic ion channel dynamics contributing to the IPSC-evoked, post-inhibitory rebound LTS. We first compared examples of LTS-*producing* responses evoked by *s*GAT1- and *s*GAT3-Block waveforms with examples of LTS-*lacking* responses evoked by *s*Control and *s*Dual-Block waveforms in a model neuron (Figure 5). For LTS-producing responses, the LTS voltage response was present across all three compartments and appeared largest in the distal dendrite (Figure 5A). This voltage response reflected a dominant T-type calcium current in dendritic compartments (Figure 5B-C), consistent with experimental findings (Munsch et al., 1997; Williams and Stuart, 2000; Zhou et al., 1997). The initiation of the LTS was also associated with a slightly delayed outward A-type potassium current that was distributed more evenly across compartments (Figure 5B-C). The temporal overlap between the T and A currents has been shown to be important for controlling the LTS amplitude and width (Pape et al., 1994).

**Figure 5.**
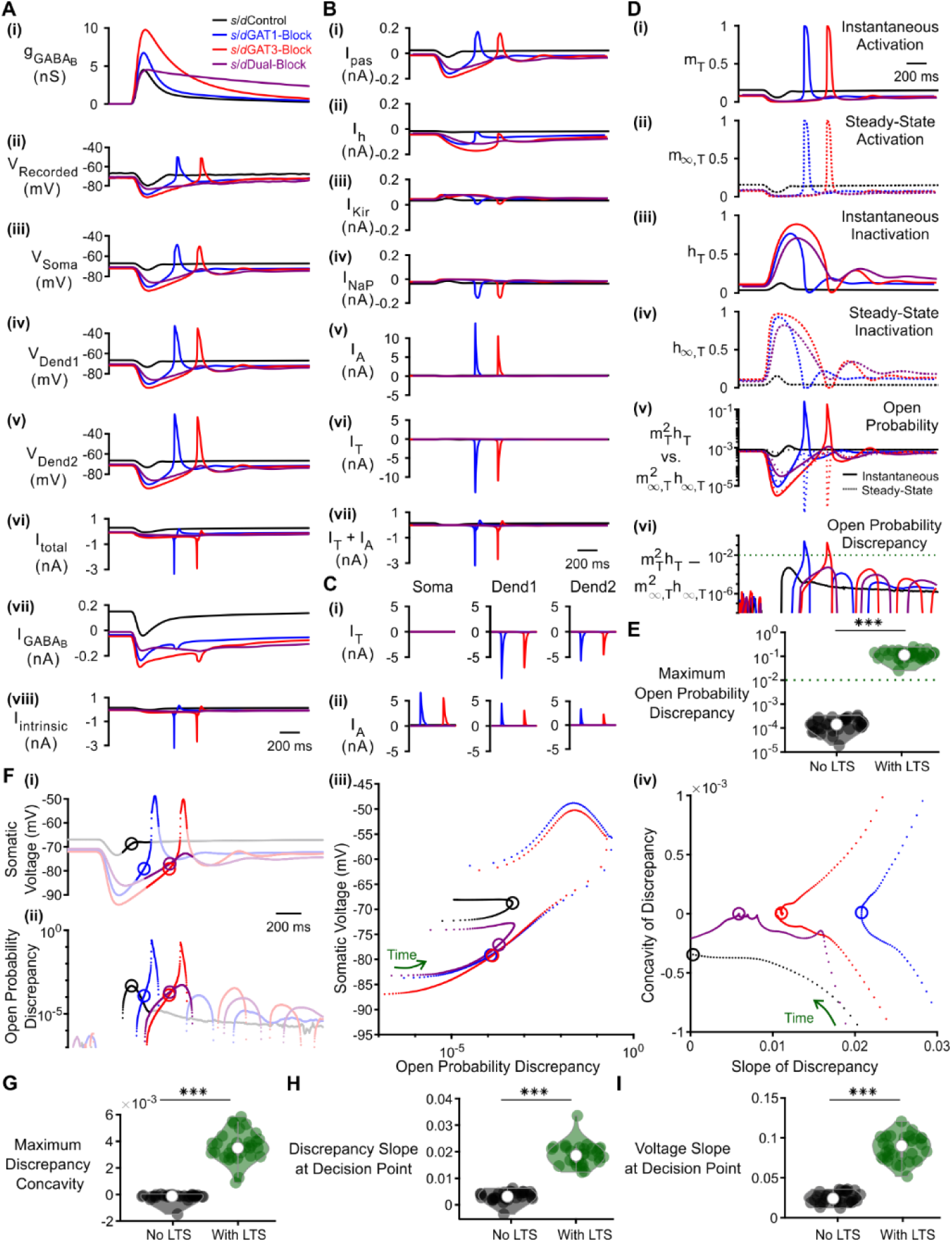
T-type calcium channel inactivation lag and open probability discrepancy depend on GABA_B_ IPSC waveforms. (**A**) The LTS response was most pronounced in the distal dendrite and correlates with the presence of large intrinsic channel currents. (**i**) Command *d*/*s*IPSCs as in Figure 2B. (**ii**) Voltage responses of Neuron 1 of Figure 2C recorded using dynamic clamp. (**iii-viii**) Simulated responses of the corresponding model neuron, including: (**iii**) somatic voltage, (**iv**) proximal dendritic voltage, (**v**) distal dendritic voltage, (**vi**) total current, (**vii**) GABA_B_ receptor current, (**viii**) total intrinsic channel current. Currents were summed over all three compartments. (**B**) T-type calcium currents and A-type potassium currents were at least an order of magnitude larger than other intrinsic currents, with the T currents contributing to LTS initiation. Intrinsic currents were summed over all three compartments and include: (**i**) passive leak current, (**ii**) hyperpolarization-activated cationic current, (**iii**) inward-rectifying potassium current, (**iv**) persistent sodium current, (**v**) A-type fast-transient potassium current, (**vi**) T-type calcium current. A sum of A and T currents is shown in (**vii**). (**C**) Comparison of A and T currents across compartments. T currents were much larger in the dendrites and A currents were slightly elevated in the soma. (**D**) A combination of T channel recovery (high *h*_*T*_) and inactivation lag (*h*_*T*_ different from *h*_*T*,∞_) appeared to be necessary for T channel opening (high *m*_*T*_ ^2^*h*_*T*_). State variables for the distal dendritic T channel (other two compartments are similar) include: (**i**) instantaneous activation gating variable, (**ii**) steady-state activation gating variable, (**iii**) instantaneous inactivation gating variable, (**iv**) steady-state inactivation gating variable, (**v**) instantaneous open probability (solid line) versus steady-state open probability (dotted line), (**vi**) difference of instantaneous versus steady-state open probability (open probability discrepancy). (**E**) The maximum open probability discrepancy was higher in LTS-producing responses (*** p < 0.001, n = 31 cells, paired-sample *t*-test). (**F**) Within the LTS regions highlighted in both (**i**) somatic voltage curves and (**ii**) dendritic T channel open probability discrepancy curves, LTS-producing responses followed different trajectories than LTS-lacking responses in either (**iii**) the voltage vs. open probability discrepancy phase plot or (**iv**) the concavity versus slope of discrepancy phase plot. Sample points plotted are 1 ms apart. Circles denote the decision points where either zero (or maximal negative) concavity is reached in the open probability discrepancy curves. (**G**) The maximum open probability discrepancy concavity in the LTS region, (**H**) The slope of open probability discrepancy at the decision point, and (**I**) The slope of somatic voltage at the decision point were all higher in LTS-producing responses (*** p < 0.001, n = 31 cells, paired-sample *t*-test).

While much is known regarding the gating properties of low threshold, T-type calcium channels, little is known about how these properties behave during physiological stimuli such as voltage changes induced by synaptic inhibition. Although it has been proposed from artificial voltage ramp studies that rebound bursting is sensitive to the slope of voltage repolarization (Gutierrez et al., 2001), the underlying T channel dynamics that confer such voltage sensitivity remain unknown. We therefore sought to understand how T channel activation and inactivation contribute to the production of the post-inhibitory rebound LTS (Figure 5D). In our well-fitted model neurons, we tracked the activation (*m*_*T*_) and inactivation (*h*_*T*_) gating variables of the T channel, as a function of time. By convention, *m*_*T*_ = 1 when all channels are activated, and *h*_*T*_ = 0 when all channels are inactivated (Hodgkin and Huxley, 1952). We also distinguished between steady-state values (*m*_*T*,∞_ and *h*_*T*,∞_), which depend only on voltage, from instantaneous values (*m*_*T*_ and *h*_*T*_), which reach steady-state values exponentially through a voltage-dependent time constant (i.e. depend on voltage and time).

One notable feature of the T channel is that the inactivation time constant is about 10-fold higher than the activation time constant (Coulter et al., 1989). Indeed, in all conditions, the instantaneous T channel activation variable *m*_*T*_ was nearly identical to its voltage-dependent steady-state value *m*_*T*,∞_. However, the instantaneous T channel inactivation variable *h*_*T*_ never achieved its steady-state value *h*_*T*,∞_ during dynamic changes in membrane voltage (Figure 5D). That is, the activation gate responded to changes in membrane potential quickly, but the inactivation gate responds slowly. Notably, the T channel inactivation lag was larger for LTS-producing responses than for LTS-lacking responses.

We next explored how T channel inactivation lag affects open probability. Without significant T channel inactivation lag, there was no discrepancy between the instantaneous and steady state T channel open probabilities. Based on the measurements by Huguenard and McCormick (Huguenard and McCormick, 1992), the maximum achievable T channel open probability at steady state (max_*V*_ (*m*_*T*,∞_(*V*)^2^*h*_*T*,∞_(*V*))) is 8.4 × 10^−4^, which is close to the LTS-lacking, pre-IPSC, baseline values in our simulations (Figure 5D). Consequently, an LTS was only produced when the instantaneous T channel open probability (*m*_*T*_^2^*h*_*T*_) was orders of magnitude higher than steady-state open probabilities. In fact, when all 31 well-fitted model neurons were considered, the maximum difference between the instantaneous and steady-state open probability (max*t* (*mT*2*hT* – *m*_*T*,∞_^2^*h*_*T*,∞_)) was on average 2.9 orders of magnitude higher for LTS-producing responses than for LTS-lacking responses (Figure 5E, n = 31 cells, p = 2.4 × 10^−29^). Herein, we refer to the difference between instantaneous versus steady-state open probability (*m*_*T*_^2^*h*_*T*_ – *m*_*T*,∞_^2^*h*_*T*,∞_) simply as *T channel open probability discrepancy*. When each trace for all 31 model neurons was considered, a threshold open probability discrepancy of 10^−2^ separated LTS-producing responses from LTS-lacking responses (not shown).

We sought to understand how high or low T channel open probability discrepancy arises in response to distinct *s*IPSCs. When the somatic voltage was plotted against the dendritic T channel open probability discrepancy, LTS-producing responses produced trajectories that were qualitatively different from the LTS-lacking responses (Figure 5F). For all *s*IPSC responses, open probability discrepancy increases upon voltage depolarization. Nevertheless, only for LTS-producing responses does the open probability discrepancy curve reach an inflection point (point of zero concavity) that eventually progresses to positive concavity. We define the point at which the open probability discrepancy curve reaches zero (or maximal negative) concavity as the *decision point* (circles in Figure 5F). A comparison between the *s*GAT3-Block (red) and sDual-Block (purple) responses showed that at the decision point, the *slope* of open probability discrepancy, rather than its value, determines whether positive concavity is eventually achieved. In fact, for all open probability curves that reach zero concavity, the slope of the open probability discrepancy curve at the decision point is always higher for LTS-producing responses than for LTS-lacking responses, but the threshold is cell-dependent (not shown). We show the progression of trajectories aligned to the decision points for two conditions (Movie 1, *s*GAT3-Block and sDual-Block) and for all conditions (Movie 2). When all 31 well-fitted model neurons were considered, there was a significant difference between LTS-producing and LTS-lacking responses for either the maximum open probability discrepancy concavity (Figure 5G), the discrepancy slope at the decision point (Figure 5H) or the voltage slope at the decision point (Figure 5I). We conclude that IPSC responses produce LTSs only if two conditions are satisfied: (1) the open probability discrepancy curve reaches zero concavity, and (2) the slope of the open probability discrepancy curve at that decision point reaches a cell-dependent threshold.

We next applied our understanding of the T channel open probability discrepancy to how GAT-modulated, GABA_B_ IPSCs regulate LTS production. We observed that the *s*Control waveform did not produce an LTS response simply because inhibition was insufficient for T channel recovery (*h*_*T*_ was always below 0.2, Figure 5D). In contrast, although T channels were sufficiently recovered (*h*_*T*_ reached above 0.6) by strong hyperpolarization associated with *s*GAT1-, *s*GAT3-, and *s*Dual-Block waveforms, only the former two waveforms produced an LTS response. We observed that rapid repolarization from a hyperpolarized state in response to the *s*GAT1- and *s*GAT3-Block waveforms allowed activation gates to open (*m*_*T*_ increased) before the inactivation gates closed (*h*_*T*_ decreased), creating a brief window characterized by a discrepancy between high instantaneous T channel open probabilities (*m*_*T*_^2^*h*_*T*_ > 10^−2^) and the low steady-state open probabilities (*m*_*T*,∞_^2^*h*_*T*,∞_ < 10^−2^). In contrast, the *s*Dual-Block waveform produced a prolonged inhibition, resulting in a slower rise during membrane potential repolarization, a small lag in T channel inactivation, a lack of increase in T channel open probability discrepancy and, ultimately, a lack of LTS response.

To test the contribution of T channel inactivation lag to LTS production, we bidirectionally altered the T channel inactivation time constant (*τ*_*h*_*T*) for the same *s*IPSC response simulations as in Figure 5. When *τ*_*h*_*T* was halved in *s*GAT1- and *s*GAT3-Block simulations, T channel open probability discrepancy remained low and LTS responses normally observed during GAT1 and GAT3 blockade were abolished (cf. Figure 6A with Figure 5). In contrast, doubling *τ*_*h*_*T* in *s*Dual-Block simulations promoted a discrepancy between instantaneous and steady-state open probabilities and, consequently, normally absent LTSs appeared (cf. Figure 6B with Figure 5). Note that *s*Control waveforms did not produce an LTS as voltage hyperpolarization was weak and T channel recovery was low. Thus, a combination of sufficient T channel recovery and high T channel open probability discrepancy appears to be necessary for LTS production.

**Figure 6.**
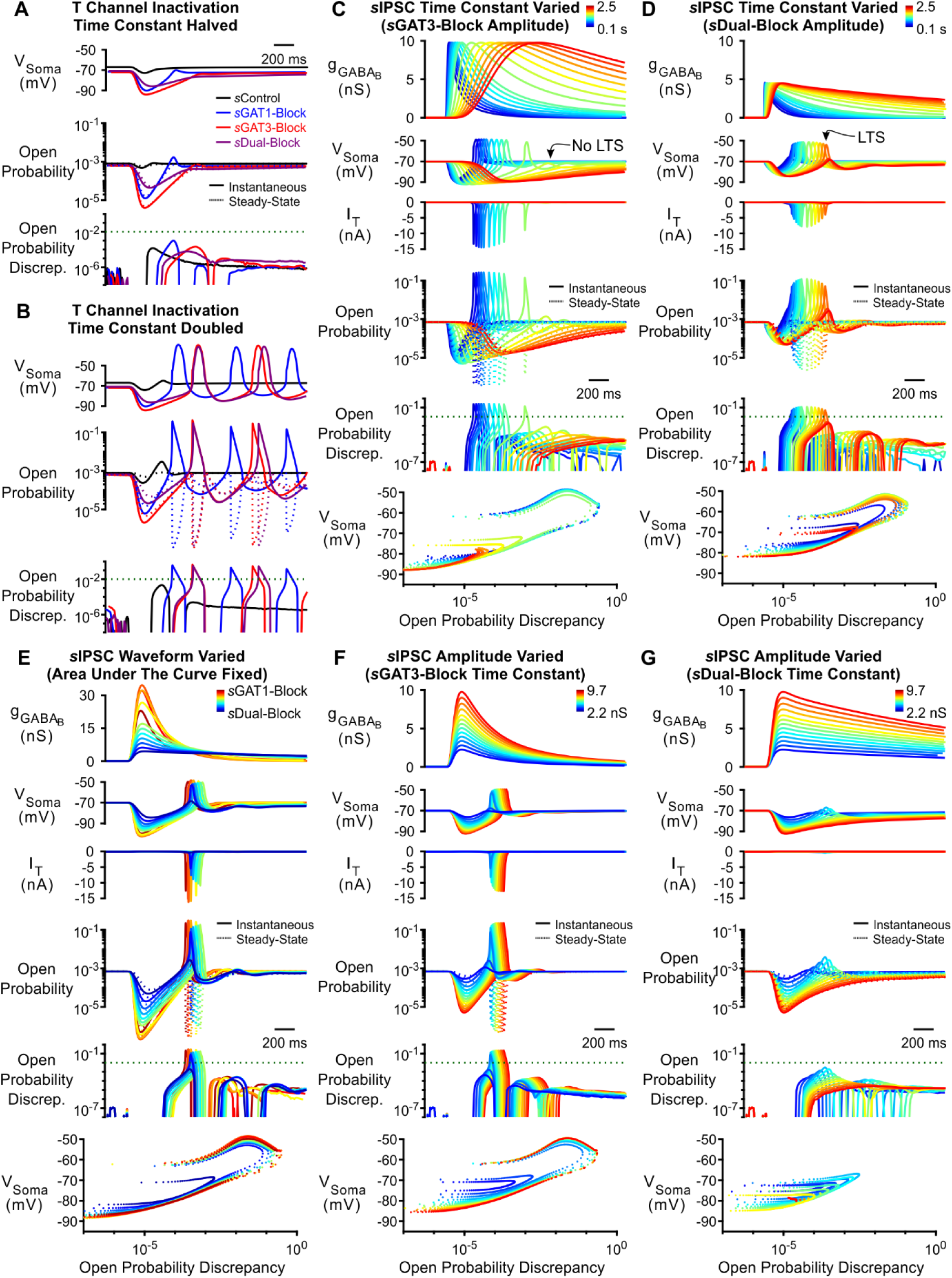
Manipulation of either the T-type calcium channel inactivation time constant or the GABA_B_ IPSC kinetics bidirectionally modulates low-threshold rebound spike production. (**A**) Simulated responses as in Figure 5 with the T channel inactivation time constant 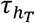 halved. The T channel inactivation lag disappeared and no LTS was produced following any *s*IPSC waveform. (**B**) Simulated responses as in Figure 5 with 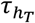 doubled. The T channel inactivation lag lengthened and LTSs appeared following *s*Dual-Block. (**C**) LTS responses to the *s*GAT3-Block waveform (blue) gradually disappeared as the time constant was increased (with amplitude fixed) to that of the *s*Dual-Block waveform (red). (**D**) LTS responses to the *s*Dual-Block waveform (red) gradually appeared as the time constant of the waveform was decreased (with amplitude fixed) to that of the *s*GAT3-Block waveform (blue). (**E**) LTS responses to the *s*Dual-Block waveform (blue) gradually appeared as all parameters of the waveform were shifted (with area under the curve fixed) to that of the *s*GAT3-Block waveform (yellow), then to the *s*GAT1-Block waveform (dark red). (**F**) LTS responses gradually appeared as the amplitude of the *s*GAT3-Block waveform was increased (with time constant fixed). (**G**) Robust LTS responses never appeared as the amplitude of the *s*Dual-Block waveform is increased (with time constant fixed).

The kinetics of inhibition appeared to underlie the observed T channel open probability discrepancy. We therefore systematically varied the time constant of an LTS-producing *s*GAT3-Block IPSC while fixing the amplitude. Prolonging IPSC kinetics decreased T channel open probability discrepancy and abolished LTS responses as time constants increased above 4-fold (Figure 6C). Conversely, shortening the time constant of a non-LTS-producing *s*Dual-Block IPSC – also while fixing the amplitude – increased T channel open probability discrepancy and produced LTS responses as time constants decreased by 20% (Figure 6D). We show the progression of trajectories aligned to the decision points for two time constants (Movie 3, last LTS success and first LTS failure) and for all time constants (Movie 4). As changing the kinetics of inhibition also changes the total amount of inhibition delivered to a cell, we also changed inhibition kinetics while keeping charge (i.e. the area under the curve) constant. Nevertheless, we continued to observe that instantaneous versus steady-state open probability discrepancies became smaller and LTS responses diminished as *s*IPSC kinetics increased (Figure 6E). Therefore, the temporal envelope of inhibition appears to be important for LTS production through its influence on T channel open probability discrepancy.

Since hyperpolarization promotes T channel recovery (Coulter et al., 1989), it remains possible that a sufficiently strong hyperpolarization – regardless of waveform – will produce an LTS. To test this possibility, we varied the inhibition amplitude using either the *s*GAT3-Block (fast kinetics) or *s*Dual-Block (slow kinetics) waveform. As we increased the amplitude of the *s*GAT3-Block waveform while fixing the rise and decay time constants, LTSs emerged as T channel open probability discrepancy increased (Figure 6F). In contrast, as the amplitude of the *s*Dual-Block waveform increased while fixing the rise and decay time constants, T channel open probability discrepancy nonetheless remained low and robust LTS responses never emerged (Figure 6G). Thus, fast inhibition kinetics are important for driving T channel open probability discrepancy, and slow kinetics provides an explanation for the consistently low LTS or burst probability across conductance amplitude scales following *s*/*d*Dual Block (Figures 2E, 4C and 4D).

In summary, LTS production following physiological inhibition is largely controlled by the dynamics of T channel open probability discrepancy in the distal dendrites. We extend this understanding by showing that LTS production appears to depend not only on the *amplitude* of inhibition, but also the *temporal envelope* of inhibition. Large inhibition amplitude is required for sufficient T channel recovery, whereas fast inhibition decay is required for driving the T channel open probability discrepancy beyond an inflection point and creating a brief time window with sufficiently high T channel open probability for LTS production.

### Network models

We next explored whether the interplay between GABA_B_-mediated inhibition and T type calcium channel dynamics in thalamocortical neurons contributes to the observed changes in network-level oscillations following GAT blockade. We first examined the effects of GABA_B_ receptor-mediated inhibition in a simplified 2-cell network configuration. In each 2-cell network, we connected a single compartment, GABAergic reticular thalamic model neuron (Klein et al., 2018) to one of the 31 well-fitted thalamocortical model neurons (Figure 7A). To generate action potentials, Hodgkin-Huxley type sodium and potassium channels were inserted into the somatic compartment of each model neuron (Williams and Stuart, 2000). The reticular thalamic neuron was connected to the thalamocortical neuron via a GABA_B_ receptor-mediated inhibitory synapse (GABA_A_ receptors were blocked during experimentally evoked oscillations, see Figure 1). Consistent with previous intra-thalamic models (Destexhe et al., 1996), the thalamocortical neuron provided AMPA receptor-mediated excitation to reticular thalamic neurons. By applying a brief stimulating current to the reticular thalamic neuron and varying the GABA_B_ receptor activation parameters (*s*IPSCs), GABA_B_ conductance waveforms comparable to those used by dynamic clamp were evoked in each thalamocortical neuron (Figure 7B). To simulate variable tonic inhibition on thalamocortical neurons, we varied the thalamocortical neuron leak reversal potential between -73 and -60 mV (14 leak reversal potentials). To generate trial-to-trial variability (5 trials per leak reversal potential), we randomized the leak conductance of each neuron to within 10% of the original value. Of all 31 possible 2-cell networks, 24 had a quiescent, pre-stimulation baseline over this range of leak reversal potentials. In those networks with quiescent baseline, oscillations persisted only when the GABA_B_-mediated inhibition promoted T channel open probability discrepancy in thalamocortical neurons to produce rebound bursting (Figure 7C). An oscillation probability was computed for each 2-cell network over the 14 × 5 = 70 trials. In addition, an oscillatory period and an oscillatory index based on the autocorrelation function of pooled spikes was computed for each successfully-evoked oscillation (see *Methods*).

**Figure 7.**
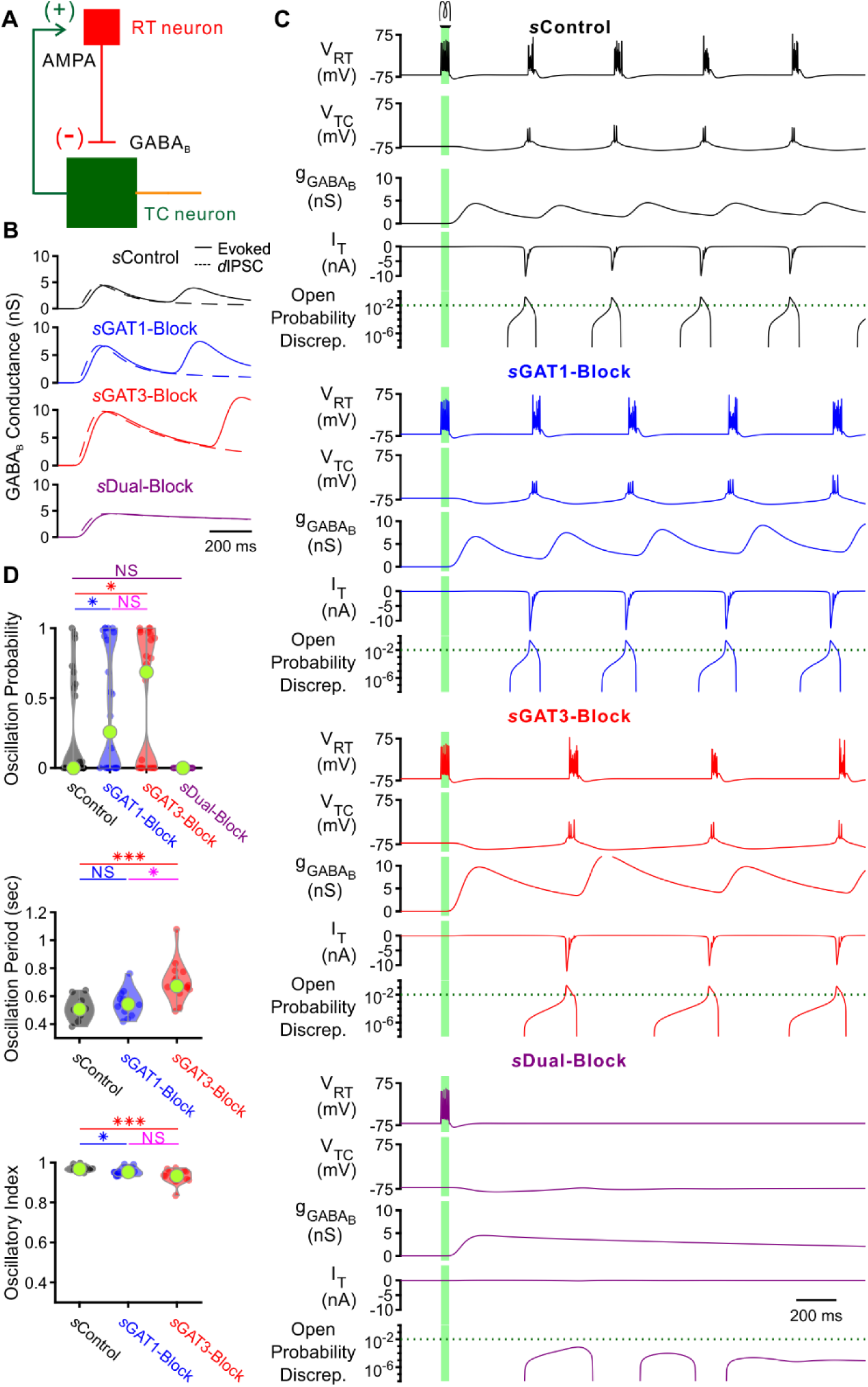
GABA_B_-receptor mediated conductance waveforms modulate oscillations produced by 2-cell model thalamic networks. (**A**) Schematic of a 2-cell model network. A reticular thalamic (RT) neuron projected a GABA_B_ receptor-mediated inhibitory synapse (-) to a thalamocortical (TC) neuron, which reciprocally projected an AMPA receptor-mediated excitatory synapse (+) to the reticular thalamic neuron. (**B**) Evoked GABA_B_ conductance waveforms in network model TC neurons (solid lines) were similar to GABA_B_ *d*IPSC waveforms (dashed lines). (**C**) 2-cell network responses under different GABA_B_ receptor conditions. Model TC neuron parameters corresponded to Neuron 1 in Figure 2. A brief (40 ms, 0.2 nA) current stimulus was applied to the reticular thalamic neuron, evoking an initial burst of 12 spikes. Oscillations were evoked under some but not all GABA_B_ receptor conditions. A total of 24 model TC neurons produced oscillations in response to stimulation. (**D**) Distributions of oscillation measures over all 24, 2-cell networks. Oscillation probability was increased when either *s*GAT1-Block parameters or *s*GAT3-Block parameters were used, but decreased when *s*Dual-Block parameters were used (* p < 0.05, ** p < 0.01, *** p < 0.001, Friedman’s test).

We examined the distributions of oscillation probability, average oscillation period and average oscillatory index over the 24 different 2-cell networks wherein *s*IPSCs were scaled by 200% (Figure 7D). Relative to using *s*Control parameters, oscillation probability increased when using *s*GAT1-Block parameters (+66%, n = 24 networks, p = 0.016) or *s*GAT3-Block parameters (+93%, p = 0.048). These results are consistent with the experimental observation that individual GAT1 or GAT3 blockade prolonged oscillations (Figure 2D). Relative to using *s*Control parameters, average oscillation period increased when using *s*GAT3-Block parameters (+39%, n = 10, p = 4.2 × 10^−4^). These results are consistent with the experimental observation that individual GAT3 blockade increased oscillation periods (Figure 2E). In contrast, when *s*Dual-Block parameters were applied in the network, oscillations do not arise, consistent with the experimental observation that dual GAT1+GAT3 blockade inevitably abolished oscillations (Figure 2D & 2F). Although the 2-cell networks recapitulated some effects of GAT blockade on oscillations, the 2-cell oscillations were extremely stereotyped and regular, resulting in unrealistically high oscillatory indices.

We sought to determine whether larger, more complex model networks could more realistically simulate experimental oscillations and recapitulate the bidirectional effects of GAT blockade by varying *s*IPSCs. We scaled up the network to include one circular layer of 100 reticular thalamic (RT) neurons and one circular layer of 100 thalamocortical (TC) neurons (Figure 8A). RT-TC inhibitory connections and TC-RT excitatory connections were both convergent and divergent. To assess the importance of the geometric and conductance heterogeneity we observed in the single cell models (Figure 3F), we established two sets of model thalamic networks: (1) 24 *TC-homogeneous* networks with TC parameters taken from each of the 24 model neurons used for the 2-cell networks and (2) 24 *TC-heterogeneous* networks with TC parameters taken from all of the 24 model neurons, randomly ordered. All networks had a quiescent, pre-stimulation baseline when the thalamocortical neuron leak reversal potential was varied between -73 and -60 mV (14 leak reversal potentials). To generate trial-to-trial variability (5 trials per leak reversal potential), we randomized the leak conductance for each of the 200 neurons to within 10% of the original value. For both *TC-homogeneous* and *TC-heterogeneous* networks, oscillations emerged and spread in response to some but not all GABA_B_ receptor activation parameters (Figure 8B & 8C). An oscillation probability was computed for each 200-cell network over the 14 × 5 = 70 trials. In addition, an oscillatory period, an oscillatory index and a half activation time was computed for each successfully-evoked oscillation (see *Methods*).

**Figure 8.**
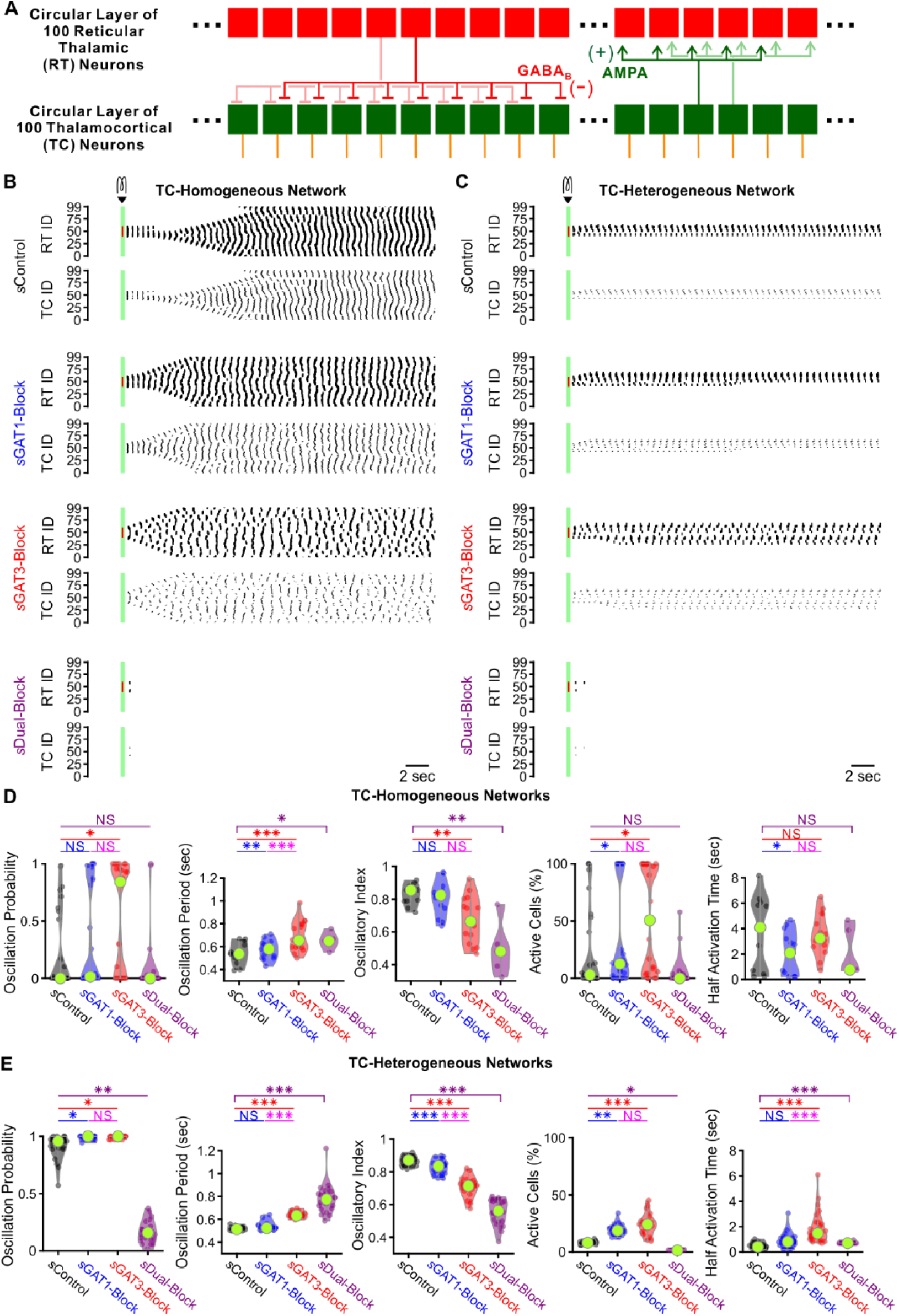
GABA_B_-receptor mediated conductance waveforms modulate oscillations produced by 200-cell model thalamic networks. (**A**) Schematic of a 200-cell model network. Each reticular thalamic (RT) neurons projected GABA_B_ receptor-mediated inhibitory synapses (-) to 9 nearby thalamocortical (TC) neurons. Each TC neuron projected AMPA receptor-mediated synapses (+) to 5 nearby reticular thalamic neurons. (**B**) Sample spike raster plots of a TC-homogenous network, using model TC neuron parameters corresponding to Neuron 1 of Figure 3. A brief (40 ms, 0.2 nA) current stimulus was applied to each of the center 20 reticular thalamic neurons. Spikes within the stimulation period are red; all other evoked spikes are black. (**C**) Sample spike raster plots of a TC-heterogeneous network, using model TC neuron parameters corresponding to the 24 model TC neurons used in Figure 7. Relative to TC-homogeneous networks, activity was more localized for TC-heterogeneous networks. (**D**) Distributions of oscillation measures over all 24 TC-homogeneous 200-cell networks (* p < 0.05, ** p < 0.01, *** p < 0.001, repeated-measures ANOVA for oscillation period and half activation time, Friedman’s test otherwise). (**E**) Distributions of oscillation measures over all 24 TC-heterogeneous 200-cell networks. Oscillation probability, oscillation period and percent of active cells increased when *s*GAT3-Block parameters were used, but decreased when *s*Dual-Block parameters were used (repeated-measures ANOVA for oscillatory index, Friedman’s test otherwise).

We examined the distributions of oscillation probability, average oscillation period, average oscillatory index, average percent of active TC cells and average half activation time over the set of *TC-homogeneous* networks (Figure 8D) and the set of *TC-heterogeneous* networks (Figure 8E) wherein *s*IPSCs were scaled by 200%. The values of oscillation periods and oscillatory indices for TC-heterogenous networks were similar to values extracted from experimental recordings (oscillation period: Figure 1E, oscillatory index: not shown). In response to the same *s*IPSC conditions, we observed highly varied (often bi-modal) oscillation responses across TC-homogeneous networks, which reflects the highly varied LTS responses across individual model TC neurons (Figure 4B). In contrast, oscillation measures were less variable across the different TC-heterogeneous networks. Therefore, cell heterogeneity averages out the LTS response variability, provides more robust network responses and accentuates the differences across *s*IPSCs. Indeed, for the set of *TC-heterogeneous* networks, relative to using *s*Control parameters, oscillation probability increased when using either *s*GAT1-Block (+8.2%, n = 24 networks, p = 0.049) or *s*GAT3-Block parameters (+8.5%, p = 0.013), but decreased when using *s*Dual-Block parameters (−82%, p = 0.0010); average oscillation period increased when using either *s*GAT3-Block (+23%, p = 1.3 × 10^−7^) or *s*Dual-Block parameters (+19%, p = 2.8 × 10^−6^); average percent of active TC cells increased when using either *s*GAT1-Block (+130%, p = 0.0044) or *s*GAT3-Block parameters (+193%, p = 1.6 × 10^−5^), but decreased when using *s*Dual-Block parameters (−79%, p = 0.037). These results are consistent with experimental findings that individual GAT1 or GAT3 blockade increased oscillation durations (Figure 2D) and oscillation periods (Figure 2E), whereas dual GAT1+GAT3 blockade eliminated oscillations (Figure 2F) in acute thalamic slices.

In summary, a population of thalamic network models was established that sufficiently recapitulated the bidirectional effects of individual versus dual GAT blockade on thalamic oscillations by merely altering the kinetics of GABA_B_-receptor inhibition. Therefore, the same interplay between GABA_B_-receptor-mediated inhibition and T channel open probability that governs thalamocortical neuron rebound bursting appears to regulate network-level oscillations. Furthermore, we found that including cell heterogeneity in network simulations provides more robust oscillations and more realistically recapitulates experimental oscillation periods by averaging out LTS response heterogeneity. Using heterogeneous networks in future studies would thus facilitate comparison across pharmacological conditions.

## Discussion

We show that seizure-like thalamic oscillations were prolonged following individual GAT1 or GAT3 blockade, yet abolished following dual GAT1+GAT3 blockade. We have also shown that, relative to control GABA_B_ IPSC waveform responses, thalamocortical neuron rebound burst probability increased following waveforms corresponding to individual GAT1 or GAT3 blockade, but decreased following waveforms corresponding to dual GAT1+GAT3 blockade. Using a population of model neurons and a population of model thalamic networks, we show that the observed thalamocortical neuron responses to GABA_B_ IPSCs and the observed oscillation changes following GAT blockade, respectively, can be recapitulated by varying the GABA_B_ receptor activation waveform. Finally, we’ve characterized a link between GABA_B_-mediated inhibition and T channel opening across both voltage and time dimensions that provides an explanation for the bidirectional effects of GAT blockade on both thalamocortical rebound bursting and seizure-like oscillations. Specifically, we identified a decision point at which the discrepancy between the instantaneous and the steady-state T channel open probability follows one of two trajectories: failing to reach a concavity and slope threshold or driving past the threshold to produce a rebound burst. These observations provide a mechanistic explanation for burst sensitivity to voltage ramps (Gutierrez et al., 2001), and burst sensitivity to both the amplitude and the decay of physiological synaptic inhibition.

### Role of thalamocortical neuron rebound bursting in generalized seizures

Thalamocortical neuron rebound bursting has long been implicated in spike-wave discharges (SWDs) observed in generalized seizures, and T channels mediate thalamocortical neuron rebound bursting (Kim et al., 2001; Porcello et al., 2003). The expression of the CaV3.1 T channel subtype by thalamocortical neurons correlates with SWD expression in both animal models (Kim et al., 2001; Broicher et al., 2008; Ernst et al., 2009) and human patients (Singh et al., 2007). Blocking T channels reduce both thalamocortical neuron bursting and oscillations in thalamic slice models (Huguenard and Prince, 1994). However, the importance of thalamocortical neuron bursting in generalized seizures remains unresolved. One recent study found reduced thalamocortical neuron firing during SWDs and a lack of seizure reduction with weak, local T channel blockade in the ventrobasal nucleus (McCafferty et al., 2018; however, strong blockade diminished SWDs), while a second recent study found that increasing thalamocortical neuron bursting increases SWDs in both epileptic mice and rats (Sorokin et al., 2017). One possibility accounting for these discrepant findings is that the population of active thalamocortical neurons during SWDs is sparse (Huguenard, 2019). Interestingly, our heterogenous network models were largely characterized by robust, yet sparse, oscillations (in contrast, homogenous network models produced oscillations that were widespread, Figure 8B and C).

Our study found a strong relationship between thalamocortical neuron bursting and epileptiform thalamic oscillations. *First*, the pharmacological conditions in which oscillation *durations* were increased (individual GAT1 or GAT3 blockade) or decreased (dual GAT1+GAT3 blockade) relative to baseline were the same conditions in which thalamocortical neuron rebound burst *probability* increased (individual GAT1- or GAT3-Block GABA_B_ *d*IPSC waveform) or decreased (Dual-Block GABA_B_ *d*IPSC waveform), relative to control *d*ISPCs. *Second*, the pharmacological conditions in which oscillation *periods* were increased (GAT1 or GAT3 blockade) relative to baseline were also the same conditions in which thalamocortical neuron rebound burst *latencies* increased (individual GAT1- or GAT3-Block *d*IPSC waveform) relative to control *d*IPSCs. *Finally*, in our 2-cell model thalamic network, each successive oscillation cycle was initiated by a thalamocortical neuron rebound burst (Figure 7C), similar to what has been reported in experiments when a reticular thalamic neuron and a thalamocortical neuron is simultaneously recorded during a thalamic oscillation (Bal et al., 1995).

### Role of the inhibitory temporal envelope in seizures

Prior experimental and computational work has suggested that a shift from GABA_A_ receptor-mediated to GABA_B_ receptor-mediated inhibition at the RT-TC synapse transforms oscillations in acute thalamic slices from a 10 Hz, sparse, spindle-like activity to a 3 Hz, hyper-synchronized, seizure-like state (von Krosigk et al., 1993; Destexhe et al., 1996; Destexhe, 1998; Blumenfeld and McCormick, 2000). Notably, the shift in oscillation frequency is consistent with differences in IPSC decay constants (GABA_A_: < 100 ms; GABA_B_: about 300 ms) recorded in thalamocortical neurons (Huguenard and Prince, 1994). Multiple animal model studies support the hypothesis that generalized spike-wave seizures rely on robust GABA_B_ receptor-mediated inhibition. Specifically, systemic injection of GABA_B_ receptor agonists increases SWDs (Liu et al., 1992; Bortolato et al., 2010), while injection of GABA_B_ receptor antagonists reduces or even abolishes SWDs (Liu et al., 1992; Vergnes et al., 1997). Notably, however, other studies were not able to record rhythmic GABA_B_ IPSCs during SWDs *in vivo* (Charpier et al., 1999) or find significant SWD changes with GABA_B_ receptor modulation (Staak and Pape, 2001), and have highlighted a particular role for extrasynaptic *GABA*_*A*_ receptors in SWDs (Cope et al., 2009). Presumably, tonic GABA_A_ currents hyperpolarize the resting membrane potential, promoting T channel recovery and increasing rebound bursting (Cope et al., 2005). Thus, the disparate conclusions regarding the importance of GABA_A_ receptor-versus GABA_B_ receptor-mediated inhibition appear to nonetheless converge on similar conclusions regarding the importance of T channel recovery, a process that involves membrane potential hyperpolarization. Under physiological conditions, it seems reasonable to expect multiple, convergent inhibitory mechanisms that promote T channel recovery.

Our study establishes a clear link between GABA_B_ receptor-mediated inhibition, thalamocortical neuron rebound bursting and epileptiform thalamic oscillations. *First*, we found a 1.4-fold or a 2-fold increase in oscillation duration in acute thalamic slices after perfusing with the GAT1 blocker NO-711 or the GAT3 blocker SNAP-5114 (Figure 1D), closely corresponding with the 1.5-fold or 2.2-fold increase in GABA_B_ IPSC amplitude recorded under the same conditions, respectively (Beenhakker and Huguenard, 2010). *Second*, in both dynamic clamp recordings and model neuron simulations, the GAT1-Block and GAT3-Block GABA_B_ IPSC waveforms produced higher thalamocortical neuron rebound LTS or burst probability relative to the Control waveform, with the GAT3-Block waveform increasing LTS or burst probability more (Figure 2D & Figure 4A-B). *Third*, we found that increasing conductances for each of the Control, GAT1-Block and GAT3-Block waveforms led to an increase in LTS or burst probability, for both dynamic clamp recordings and model neuron simulations (Figure 2E & Figure 4C-D). *Finally*, a comparison of the Control (black) versus GAT1-Block (blue) IPSC responses in Figure 5 shows that the lack of LTS in the former correlates with a decreased level of TC hyperpolarization, a lack of T channel de-inactivation (both *h*_*T*_ and *h*_∞,*T*_ are low) and a deficiency in T current production (no spike in *I*_*T*_), agreeing with prior studies in that the initiation of a low-threshold rebound spike depends on the sufficient removal of T channel inactivation through membrane potential hyperpolarization (Llinás and Jahnsen, 1982; Coulter et al., 1989).

More interestingly, we found that not only is the overall amount of inhibition important, how such inhibition distributes over time is equally important. For instance, the *d*Dual-Block waveform has an area under the curve (i.e. charge) about twice that of the *d*GAT3-Block waveform (Figure 2B). Nevertheless, both dynamic clamp recordings and model neuron simulations showed that the *d*Dual-Block waveform decreased rebound burst probability relative to the control, whereas the *d*GAT3-Block waveform increased rebound burst probability (Figure 2D & Figure 4). In fact, the *d*Dual-Block waveform largely failed to produce rebound bursts even when the conductance amplitude was scaled so high that the burst probability was close to 1 in all other conditions, i.e. following *d*Control, *d*GAT1-Block and *d*GAT3-Block waveforms (Figure 2E & Figure 4). Dual GAT blockade also abolished oscillations, in stark contrast to the robust prolongation of oscillations observed during GAT3 blockade only (Figure 1D & 1F). Therefore, even high levels of synaptic inhibition, if decayed too slowly, can abolish both thalamocortical neuron rebound bursts and epileptiform thalamic oscillations, a conclusion recapitulated by our model thalamic networks (Figure 7D & 8C).

### T channel open probability discrepancy drives thalamocortical neuron rebound bursting

What are the intrinsic channels in thalamocortical neuron that enable rebound bursting? Prior experimental work has shown that T channels are critical for the generation of post-inhibitory low-threshold rebound spikes in thalamocortical neurons (Kim et al., 2001; Porcello et al., 2003). A computational study by Destexhe *et al*. (1998) showed that membrane voltage trajectories of low-threshold rebound spikes in thalamocortical neurons can be recapitulated using a 3-compartment model, but not a single-compartment model. A recent study by Amarillo *et al*. (Amarillo et al., 2014) used pharmacological approaches to identify seven intrinsic channels (*I*_*T*_, *I*_*h*_, *I*_*A*_, *I*_*Kir*_, *I*_*NaP*_, *I*_*NaLeak*_, *I*_*KLeak*_) contributing to the resting membrane potential of thalamocortical neurons. The same study created a single-compartment computational model neuron that produced thalamocortical neuron rebound bursting and showed that a balance between the 5 voltage-dependent channels shapes the low-threshold rebound spike.

Our model thalamocortical neurons extend the studies by Destexhe et al. (1998) and Amarillo *et al*. (2014). That is, we incorporated the same intrinsic channels described by Amarillo *et al*. (2014), but allowed channel density to vary across three compartments [note that the two different leak channels in Amarillo *et al*. were simplified by combining them into a single passive leak conductance (*g*_*pas*_) with a reversal potential (*E*_*pas*_)]. Furthermore, we established a different model neuron for each set of single thalamocortical neuron recordings, in an effort to capture the heterogeneity in response to physiologically relevant inhibitory inputs such as GABA_B_ receptor-mediated IPSCs (Figure 3). By creating a set of 31 well-fitted model neurons, we were also able to generate population statistics that were in many ways similar to those of the corresponding population of recorded neurons (Figure 4).

We found that the well-fitted neurons, despite having fitted to IPSC responses that were widely heterogeneous in both LTS probability and latency, two parameters were highly convergent. *First*, T channel densities were always higher in the dendrites than in the soma (Figure 3F). This observation is in agreement with prior computational and experimental work (Munsch et al., 1997; Zhou et al., 1997; Destexhe et al., 1998; Williams and Stuart, 2000). In fact, as shown in Figure 5C, the relative magnitude of T currents during the LTS in all of our well-fitted models was negligible in the soma relative to that in the dendrites. Further, even after reducing the somatic T currents to zero, there was no visible change in the LTS responses (data not shown). Thus, our simulations suggest that the LTS response in the soma that ultimately results in a burst of action potentials is highly dependent on dendritic input currents. *Second*, A-type potassium channel densities were always highest in the soma (Figure 3F). As A channels are important for controlling the amplitude and width of the LTS response, but not for initiation (Amarillo et al., 2014), we focused on the relationship between inhibition and T channel opening in this study. Nevertheless, the striking similarity of the sum of the T and A currents (*I*_*T*_ + *I*_*A*_) versus the total currents provided by intrinsic channels (*I*_*Intrinsic*_) suggests that these two currents work in opposition to shape the LTS response (Figure 5A-B, Pape and McCormick, 1995). The degree to which modulation of the A channel influences thalamocortical neuron bursting has yet to be explored.

Perhaps more interestingly, our model neuron allowed us to identify a novel mechanism in which inhibition kinetics regulate T channel opening. We discovered a pronounced lag between the instantaneous and steady-state T channel inactivation curves (*h*_*T*_ versus *h*_*T*,∞_) during *s*GAT1-Block and *s*GAT3-Block waveforms, but not during *s*Dual-Block waveforms. As this lag was difficult to visualize and was only important when the activation (*m*_*T*_) was also high, we computed the instantaneous T channel open probability (*m*_*T*_^2^*h*_*T*_) versus its steady-state (*m*_*T*,∞_^2^*h*_*T*,∞_) and resolved a high discrepancy during *s*GAT1- and *s*GAT3-Block waveforms, but not during *s*Dual-Block waveforms. Across all GABA_B_ *s*IPSC responses, the maximum difference between instantaneous versus steady-state T channel open probability predicted the presence of an LTS response (Figure 5E). Indeed, upon bidirectional manipulation of T channel inactivation time constant, congruent, bidirectional changes in T channel open probability discrepancy and LTS production were observed (Figure 6A-B). We thus propose that future efforts in modulating thalamocortical rebound bursting can be directed at modulating the T channel inactivation time constant.

In order to confirm that inhibition kinetics influence LTS production by controlling T channel opening, we varied the GABA_B_ *s*IPSC waveform systematically five different ways (Figure 6C-G). In each case, an LTS was only produced when there was both sufficient T channel recovery and an increase in T channel open probability discrepancy. We concluded that a high inhibition amplitude was required for sufficient T channel recovery, but fast inhibition kinetics was required for driving T channel open probability discrepancy past the threshold for LTS production. Collectively, the results of these manipulations (c.f. Figures 6F and 6G) support the hypothesis that dual GAT1+GAT3 blockade eliminates thalamic oscillations by promoting sustained, GABA_B_ receptor-mediated inhibition that, in turn, promotes a convergence of instantaneous and steady-state T channel open probabilities, a state that results in LTS failure.

### Implication on anti-epileptic therapies

Drugs that increase synaptic inhibition in the brain may paradoxically exacerbate generalized seizures. For example, the GAT1 blocker tiagabine increases extracellular GABA concentrations (Beenhakker and Huguenard, 2010) and is an effective anti-epileptic drug used to treat focal epilepsy (Nielsen et al., 1991; Uthman et al., 1998). However, tiagabine can also induce continuous SWDs (i.e. absence status epilepticus) in patients with absence epilepsy (Knake et al., 1999) and non-convulsive status epilepticus in patients with focal epilepsy (Ettinger et al., 1999; Vinton et al., 2005). GAT1 knockout mice, which have increased inhibitory currents, and presumed increased activation of GABA_B_ receptors (Beenhakker and Huguenard, 2010) in thalamocortical neurons, also exhibit increased incidence of SWDs (Cope et al., 2009). Furthermore, application of GABA_B_ receptor antagonists increases seizures in rats susceptible to convulsive focal seizures, but suppresses seizures in rats with non-convulsive absence seizures (Vergnes et al., 1997). Together, these observations suggest that, although a seizure is a manifestation of hyperexcitable neuronal firing, increasing inhibition in the form of neuronal hyperpolarization may not always reduce seizures (Beenhakker and Huguenard, 2009).

In fact, as we discovered in this study, the temporal envelope of inhibition seems to play an important role in determining whether thalamocortical rebound bursts and seizure-like oscillations occur. Although the finding that dual GAT1+GAT3 blockade abolished epileptiform oscillations may at first glance suggest that nonspecific GABA transporter blockers may be used to treat generalized seizures, this approach would be relatively non-specific, including for example, disrupting normal spindle oscillation formation, leading to undesirable side effects. Nevertheless, it is possible that temporary, pharmacological enhancement of GABA transporter expression or function may create pockets of IPSC response heterogeneity in the overall thalamocortical neuron population, making it less likely for the thalamic network to develop a hypersynchronous state (Pita-Almenar et al., 2014).

## Materials and Methods

### Oscillation recordings in acute thalamic slices

Sprague-Dawley rats (Charles River Laboratories, Wilmington, MA) of postnatal day 11 to 17 (P11-P17), of either sex, were used in oscillation experiments, which were performed in accordance with protocols approved by the Institutional Animal Care and Use Committee at the University of Virginia (Charlottesville, VA). Rats were deeply anesthetized with pentobarbital, then transcardially perfused with ice-cold protective recovery solution containing the following (in mM): 92 NMDG, 25 glucose, 5 Na-ascorbate, 3 Na-pyruvate, 2.5 KCl, 2 thiourea, 20 HEPES, 26 NaHCO_3_, 1.25 NaH_2_PO_4_, 0.5 CaCl_2_, 10 MgSO_4_, titrated to a pH of 7.3-7.4 with HCl (Ting et al., 2014). Horizontal slices (400 μm) containing the thalamus were cut in ice-cold protective recovery solution using a vibratome (VT1200, Leica Biosystems, Wetzlar, Germany). Slices were trimmed to remove the hippocampus and the hypothalamus, and then transferred to protective recovery solution maintained at 32-34°C for 12 min. Brain slices were kept in room temperature ACSF consisting of the following (in mM): 126 NaCl, 2.5 KCl, 10 glucose, 26 NaHCO_3_, 1.25 NaH_2_PO_4_, 2 CaCl_2_, and 1 MgSO_4_. All solutions were equilibrated with 95% O_2_/5% CO_2_.

Slices were placed in a humidified, oxygenated interface recording chamber and perfused with oxygenated ACSF (2 mL/min) at 32-34°C. 10 μM bicuculline was added to the ACSF (bicuculline-ACSF) to block GABA_A_ receptors. Oscillations were evoked by a square voltage pulse (10 V, 0.5 ms duration) delivered once every 60 seconds through two parallel tungsten electrodes (50-100 kΩ, FHC) 50-100 μm apart and placed in either the internal capsule or the reticular thalamus, which stimulated traversing corticothalamic and thalamocortical axons. Extracellular potentials were recorded in a differential manner with two tungsten electrodes (50-100 kΩ, FHC) by placing one in the somatosensory ventrobasal nuclei of the thalamus close to the stimulating electrode and one far away from the stimulating electrode. One experiment was performed per slice. Multi-unit recordings were amplified 10,000 times with a P511 AC amplifier (Grass), digitized at 10 kHz with Digidata 1440A, band-pass filtered between 100 Hz and 3kHz, and acquired using Clampex 10.7 software (Molecular Devices, San Jose, CA).

After at least 20 minutes in baseline bicuculline-ACSF, slices were perfused with one of 4 possible solutions: (1) bicuculline-ACSF, (2) bicuculline-ACSF plus 4 μM NO-711 to block GAT1 transport (Sitte et al., 2002), (3) bicuculline-ACSF plus 100 μM SNAP-5114 to block GAT3 transport (Borden et al., 1994) or (4) bicuculline-ACSF plus a combination of 4 μM NO-711 and 100 μM SNAP-5114 to simultaneously block both GAT1 and GAT3. After 40 minutes, the perfusion solution was switched back to bicuculline-ACSF for at least 20 minutes.

### Dynamic clamp recordings

Male Sprague-Dawley rats of postnatal day 11 to 15 (P11-P15) were used in dynamic clamp experiments, which were performed in accordance with protocols approved by the Administrative Panel on Laboratory Animal Care at Stanford University. Rats were deeply anesthetized with pentobarbital, then brains were rapidly extracted and placed in ice-cold protective recovery solution containing the following (in mM): 34 sucrose, 2.5 KCl, 11 glucose, 26 NaHCO_3_, 1.25 NaH_2_PO_4_, 0.5 CaCl_2_, and 10 MgSO_4_, titrated to a pH of 7.4 with HCl. Horizontal slices (300 μm) containing the thalamus were cut in ice-cold protective recovery solution using a vibratome (VT1200, Leica Biosystems). Slices were transferred to artificial cerebrospinal fluid (ACSF) maintained at 32°C for 45-60 min, then gradually brought to room temperature. The ACSF contained the following (in mM): 126 NaCl, 2.5 KCl, 10 glucose, 26 NaHCO_3_, 1.25 NaH_2_PO_4_, 2 CaCl_2_, and 1 MgSO_4_. All solutions were equilibrated with 95% O_2_/5% CO_2_.

Slices were placed in a submerged recording chamber and perfused with oxygenated ACSF (2 mL/min) at 32-34°C. The chamber contained nylon netting which suspended the slice 1-2 mm from the chamber floor and enhanced slice perfusion. Slices were visualized with Dodt-contrast optics (Luigs & Newmann, Ratingen, Germany) on an Axioskop microscope (Zeiss, Pleasanton, CA). Recordings were obtained with a MultiClamp 700A patch amplifier (Molecular Devices), digitized with Digidata 1322A and acquired using Clampex software. Borosilicate glass pipettes (1.5-3 MΩ) pulled on a P-87 micropipette puller (Sutter Instruments, Novato, CA) were filled with an internal solution containing (in mM): 100 potassium gluconate, 13 KCl, 10 EGTA, 10 HEPES, 9 MgCl_2_, 2 Na_2_-ATP, 0.5 Na-GTP, 0.07 CaCl_2_ (pH 7.4).

Dynamic clamp experiments were conducted using a computer running the RealTime Application Interface for Linux (RTAI, www.rtai.org) sampling the intracellular potential at 50 kHz. Custom-written software modified from previous work (Sohal et al., 2006) used the sampled membrane potential and the pre-specified GABA_B_-mediated inhibitory conductance to, at every timestep, update the inhibitory current that was injected into the thalamocortical neuron at 50 kHz. All measured voltages were corrected for a junction potential of -10 mV.

Inhibitory conductance waveforms (*d*Control, *d*GAT1-Block, *d*GAT3-Block, *d*Dual-Block, Figure 2B) for use in dynamic clamp experiments were created based on previous recordings of GABA_B_-mediated IPSCs (Beenhakker and Huguenard, 2010). In the previous study, voltage-clamped thalamocortical neurons were recorded during electrical stimulation of presynaptic reticular thalamic neurons while the thalamic slices were bathed in ionotropic glutamate receptor and GABA_A_ antagonists to isolate GABA_B_-mediated IPSCs. After acquiring baseline data, one of three drugs was added to the perfusing ACSF: 4 μM NO-711 to block GAT1, 100 μM SNAP-5114 to block GAT3, or both 4 μM NO-711 and 100 μM SNAP-5114 to block both GAT1 and GAT3 (Borden et al., 1994; Sitte et al., 2002). After at least 10 minutes of drug perfusion and stabilization of drug effect, a series of GABA_B_-mediated IPSCs was acquired. In the present study, we converted the previously obtained averaged IPSCs for each drug condition into a conductance waveform fitted with the equation:

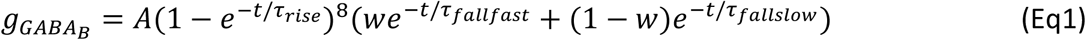

where *A* is the amplitude coefficient, *τ*_*rise*_ is the rise time constant, *τ*_*fallfast*_ and *τ*_*fallslow*_ are the fast and slow decay constants, respectively, and *w* is the weighting factor for the two decay terms (Otis et al., 1993). The percent change in each parameter from baseline to drug condition was computed for each experiment. Baseline-normalized parameter values were compared between conditions, and for parameters with significant population differences in values (p < 0.05, Wilcoxon signed rank test), the mean change in parameter value was found and implemented for that drug condition. For parameters that did not change significantly in the drug condition compared to baseline, the parameter value was kept identical to that of the baseline (*d*Control) condition for the templates (Figure 2B). Parameter values for the four experiment conditions (*d*Control, *d*GAT1-Block, *d*GAT3-Block, *d*Dual-Block) are listed in Table 1. A value of -115 mV was used to for the reversal potential associated with the GIRK conductance following post-synaptic GABA_B_ receptor activation.

**Table 1.** Parameter values of GABA_B_-mediated inhibitory conductance waveforms used in dynamic clamp experiments. These values (with the amplitudes scaled by 200%) correspond to the conductance templates shown in Figure 2B. *A*, amplitude coefficient; *τ*_*rise*_, rise time constant; *τ*_*fallfast*_, fast decay time constant; *τ*_*fallslow*_, slow decay time constant; *w*, weighting factor for the two decay terms.

| | $A$ | $\tau_{rise}$ | $\tau_{fallfast}$ | $\tau_{fallslow}$ | $w$ |
| --- | --- | --- | --- | --- | --- |
| $dControl$ | 16.00 | 52.00 | 90.10 | 1073.20 | 0.952 |
| $dGAT1$-Block | 24.00 | 52.00 | 90.10 | 1073.20 | 0.952 |
| $dGAT3$-Block | 8.88 | 38.63 | 273.40 | 1022.00 | 0.775 |
| $dDual$-Block | 6.32 | 39.88 | 65.80 | 2600.00 | 0.629 |

Since the number of GABA_B_ receptors activated in a physiological setting was not determined, all GABA_B_ IPSC waveforms (*d*IPSCs) had amplitude scaled between 25%-800%. A total of 5-15 repetitions were performed for each *d*IPSC waveform, with a holding current adjusted at the beginning of each recording so the holding membrane potential is in the range of -73 to -60 mV. For each sweep, a brief current pulse (−50 pA, 10 ms) was applied at around 100 ms to assess electrode resistance and passive membrane properties, then the *d*IPSC was applied at 1000 ms.

### Analysis of oscillation recordings

MATLAB R2018a (MathWorks, Natick, MA) was used for all data analysis. Single action potential *spikes* were detected from raw multi-unit activity as follows (Sohal et al., 2003). The raw signal was first bandpass-filtered between 100-1000 Hz. Slopes between consecutive sample points were computed from the filtered signal. For each sweep *i*, the baseline slope noise *α*_*i*_ was computed from the root-mean-square average of the slope vector over the baseline region before stimulation start, and the maximum slope *β*_*i*_ was defined as the maximum slope value at least 25 ms after stimulation start (to account for the stimulus artifact). For each slice, if *α* = ∑_*i*_ *α*_*i*_ is the baseline slope noise averaged over all sweeps and *β* = ∑_*i*_ *β*_*i*_ is the maximum slope averaged over all sweeps, then the slice-dependent signal-to-noise ratio was given by 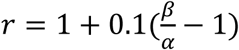. The resulting signal-to-noise ratios fell between 2-5. A sweep-dependent slope threshold was then defined by *θ*_*i*_ = *rα*_*i*_. Finally, single spikes were defined as all local maxima of the slope vector at least 25 ms after stimulation start with values exceeding a slope threshold.

To compute the oscillation duration for each sweep, spikes were first binned by 10 ms intervals to yield a spike histogram. Evoked *bursts* were detected by joining consecutive bins with a minimum spike rate of 100 Hz, using a minimum burst length of 60 ms, a maximum delay after stimulation start of 2000 ms and a maximum inter-burst interval of 2000 ms (Figure 1A). The *oscillation duration* was defined as the time difference between the end of the last evoked burst and stimulation start.

To compute the oscillation period for each sweep, an autocorrelation function (ACF) was first computed from the binned spikes, then moving-average-filtered using a 100 ms window. Peaks (local maxima) were detected from the filtered ACF using a minimum peak prominence that is 0.02 of the amplitude of the primary peak (the first value of the ACF). Peaks with lags greater than the oscillation duration were ignored. The *oscillation period* was computed by searching for the lag value *δ* that minimizes the distance function *f*(*δ*) = ∑_*j*_ |*p*_*j*_ – *q*_*j*_(*δ*)|, where *p*_*j*_ is the lag value of peak *j* and *q*_*j*_(*δ*) is the closest multiple of *δ* to *p*_*j*_. The search was initialized with the estimate *δ*0 = *p*2 – *p*1 and confined to the bounds 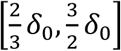.

An *oscillatory index* (Sohal et al., 2006) was also computed from the filtered autocorrelation function (fACF). For each sweep, the non-primary peak with largest fACF value was designated as the secondary peak. The *oscillatory index* was then defined by *OI* = (*A*_*peak*2_ – *A*_*trough*_)/(*A*_*peak*1_ – *A*_*trough*_), where *A*_*peak*1_ is the fACF value of the primary peak, *A*_*peak*2_ is the fACF value of the secondary peak and *A*_*trough*_ is the minimum fACF value between the primary peak and the secondary peak.

The average oscillation duration or period at the end of each phase (baseline or drug) was computed by choosing the 5 values from the last 10 sweeps of each phase that were within 40% of the average of the group of values. This approach was used to minimize bias caused by abnormally-shortened evoked oscillations due to the presence of spontaneous oscillations.

### Analysis of dynamic clamp recordings

MATLAB R2018a was used for all data analysis. Responses to *d*IPSCs were analyzed as follows: All voltage traces were manually examined and noisy recordings were excluded. The peak of the current trace (IPSC peak) within the first 300 ms of *d*IPSC start was first detected. The most likely candidate for a calcium-dependent low-threshold spike (LTS) was then detected from the raw voltage trace in between the time of IPSC peak and 7000 ms after *d*IPSC start.

To detect the *LTS candidate*, the voltage trace was first median-filtered with a time window of 30 ms to remove action potentials (Chung et al., 2002), then moving-average-filtered with a time window of 30 ms. First and second derivatives of the doubly-filtered voltage traces were computed by differences between consecutive sample points, and the first derivative vector was moving-average-filtered with a time window of 30 ms before taking the second derivative. The local maximum of the doubly-filtered voltage trace with the most negative second derivative was chosen as the LTS candidate. A histogram of LTS candidate second derivatives for traces was fitted to a sum of 3 Gaussian distributions and the minimum of the probability density function between the first two peaks was used as a second derivative threshold.

Features were computed for each LTS candidate. The LTS *peak value* was the absolute voltage value for the peak. The LTS *latency* was defined by the time difference between the LTS peak time and *d*IPSC start. A time region that was bounded by the first local minimum of the doubly-filtered voltage trace on either side of the peak was used to compute the LTS *maximum slope* and detect action potentials. For accuracy, the slope value was computed from a different doubly-filtered trace with a smaller moving-average-filter time window corresponding to roughly a 3 mV-change in amplitude for that trace. Action potentials spikes were detected by a relative amplitude threshold 10 mV above the LTS amplitude for that trace.

An LTS candidate was then assigned as an *LTS* if the following criteria were all satisfied: (1) Peak prominence must be greater than the standard deviation of the filtered voltage values before IPSC start; (2) Peak second derivative must be more negative than the threshold (−0.0023 V^2^/s^2^) described above; (3) If there were action potentials riding on the LTS, the LTS peak time must occur after the time of the first action potential. Detection results were manually examined by blinded experts and the algorithm’s decision for LTS determination was overturned only if 3 of the 4 polled electrophysiology experts agreed. The *LTS or burst probability* was defined as the proportion of traces producing an LTS or a *burst* (an LTS with at least one riding action potential) across all 5-15 repetitions (with varying holding potentials) for a particular GABA_B_ IPSC waveform.

### Single thalamocortical neuron models

All computational simulations were performed using NEURON (Carnevale and Hines, 2009) version 7.5 with a temperature of 33°C. Each model thalamocortical neuron had one cylindrical somatic compartment (Soma) and two cylindrical dendritic compartments (Dend1 and Dend2) in series (Figure 3C). The somatic length and diameter were set to be equivalent with the parameter *diam*_*soma*_. The two dendritic compartments were set to have equal diameters (*diam*_*dend*_), with each having lengths equal to half of the parameter *L*_*dend*_. Values for specific membrane capacitance and axial resistivity were equivalent to those reported by Destexhe et al. (1998). Passive leak channels were inserted in all three compartments at equivalent densities (*g*_*pas*_) with a fixed reversal potential of -70 mV. The 4 voltage-independent (*passive*) parameters described above were allowed to vary across neurons.

The following mechanisms were inserted in all 3 compartments: the T-type calcium current (*I*_*T*_) and the submembranal calcium extrusion mechanism (*Ca*_*decay*_) was adapted from Destexhe et al. (1998); the hyperpolarization-activated nonspecific cationic current (*I*_*h*_), the A-type transient potassium current (*I*_*A*_), the inward-rectifying potassium current (*I*_*Kir*_) and the persistent sodium current (*I*_*NaP*_) were adapted from Amarillo et al. (2014). The following parameters were allowed to vary across compartments and across neurons: the maximum permeability 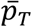 (in cm/s) of *I*_*T*_, which was described by the Goldman–Hodgkin–Katz flux equation (Huguenard and McCormick, 1992), and the maximum conductance densities 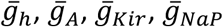 (in S/cm^2^) of other currents described by Ohm’s law. All parameters that were not varied during optimization were identical for all model neurons and taken from literature values. Based on calculations from solutions used in experiments, the potassium reversal potential was -100 mV, the sodium reversal potential was 88 mV, the calcium concentration outside neurons was 2 mM and the initial calcium concentration inside neurons was 240 nM.

All intrinsic current mechanisms were described by Hodgkin-Huxley type equations with voltage and/or time-dependent activation (*m*) and inactivation (*h*) gating variables and have been described in detail in the corresponding sources (Amarillo et al., 2014; Destexhe et al., 1998). In particular, the open probability of *I*_*T*_ was described by *m*(*V, t*)^2^*h*(*V, t*) (Huguenard and Prince, 1992). At every point in time, the activation variable *m* converges exponentially to its steady-state value *m*_∞_(*V*) with a time constant *τ*_*m*_(*V*) and the inactivation variable *h* converges exponentially to its steady-state value *h*_∞_(*V*) with a time constant *τ*_*h*_(*V*). The equations for *m*_∞_(*V*), *τ*_*m*_(*V*), *h*_∞_(*V*) and *τ*_*h*_(*V*) were fitted from electrophysiological recordings and described by Huguenard & McCormick (1992).

For simulations involving action potentials, a fast sodium and potassium mechanism adapted from Sohal and Huguenard (2003) was inserted in the somatic compartment and made identical across neurons. Based on the comparison of the number of spikes per LTS between model neurons and corresponding recorded neurons (Figure 4A-B), the threshold parameter *V*_*Traub*_ was changed to -65 mV.

A custom GABA_B_ receptor mechanism was inserted in the somatic compartment that produced IPSCs with the equation form given by Eq1. The parameters used were identical to that used for dynamic clamp, with kinetics varying for each pharmacological condition as in Table 1. Although parameter measurements were performed at a same temperature (33 °C) as simulations, a Q_10_ of 2.1 (Otis et al., 1993) was included in the model.

All simulations performed for each model neuron were matched to traces recorded using dynamic clamp for the corresponding recorded neuron as follows: To match the holding potentials, we first performed a test simulation to voltage clamp the model neuron (with initial potential -70 mV) for 2 seconds to bring it to a quasi-steady state. The resultant steady-state current was used as holding for the current clamp simulation. To simulate dynamic clamp of the recorded neuron, after 2 seconds to allow state variables in the model to stabilize, a brief current pulse (−50 pA, 10 ms) was applied at around 2.1 seconds, then an *s*IPSC identical to the *d*IPSC applied in dynamic clamp was applied at 3 seconds. The analysis of LTS and burst features for simulated *s*IPSC responses is identical to that for recorded *d*IPSC responses.

### Parameter initialization

Initial values for *passive parameters* were different for each model neuron and were estimated from the current pulse responses from the corresponding recorded neuron by a strategy adopted from Johnston and Wu (Johnston and Wu, 1994). A portion of raw current pulse responses had a systematic voltage shift at the beginning and end of the pulse, consistent with an unbalanced bridge. These shifts were detected by a slope threshold determined through a histogram of all initial slopes and then corrected by shifting the entire portion of the response during the brief current pulse, by the calculated amount. The corrected current pulse responses for a particular neuron were then fitted to the following first order response equation with 2 exponential components:

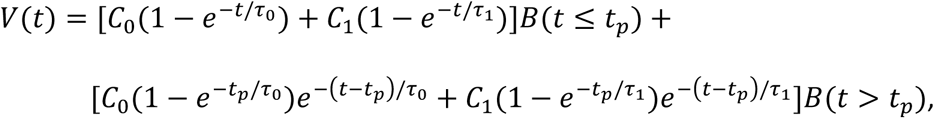

where *C*_0_, *C*_1_ are the amplitudes (in mV) of the two components, *τ*_0_, *τ*_1_ are the time constants (in ms) of the two components, *t*_*p*_ = 10 ms is the width of the current pulse, and *B*(*x*) is a Boolean function defined by *B*(*x*) = 1 if *x* = true and *B*(*x*) = 0 if *x* = false. Initial values for the curve fit were *C*_0,*i*_ = Δ*V, C*_1,*i*_ = Δ*V, τ*_0,*i*_ = 10 ms, *τ*_1,*i*_ = 1 ms where Δ*V* is the mean voltage change for the neuron after each 10 ms stimulus.

Next, given the fitted coefficients, we estimated the defining parameters for a ball-and-stick model. The input resistance was computed by

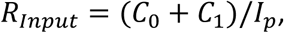

where *I*_*p*_ = 50 pA is the amplitude of the current pulse. The membrane time constant *τ*_*m*_ was set to equal that of the slow component *τ*_*m*_ = *τ*_0_. The length constant was computed by solving for *L* in equation 4.5.57 of Johnston and Wu:

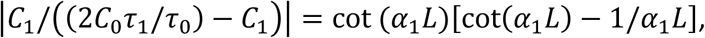

where *α*_1_ = ((*τ*_1_/*τ*_0_)−1)^1/2^, starting with the initial guess of *L*_*i*_ = *Π*/*α*_1_. The dendritic-to-somatic conductance ratio *ρ* was then computed by equation 4.5.58 of Johnston and Wu:

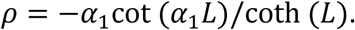

Finally, we converted the defining ball-and-stick parameters to initial estimates for the passive parameters in our 3-compartment NEURON model that are optimized. The input resistances of the soma and the dendrite were computed by

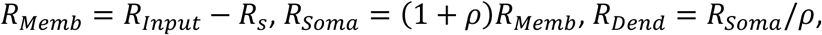

where *R*_*s*_ is the series resistance. The specific membrane resistivity *R*_*m*_ was computed by equation 4.3.3 of Johnston and Wu:

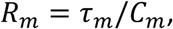

where a fixed value of 0.88 µF/cm^2^ (Destexhe et al., 1998) for the specific membrane capacitance *C*_*m*_. Then the somatic diameter *diam*_*soma*_ was computed by equation 4.3.8 of Johnston and Wu:

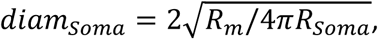

the dendritic diameter *diam*_*Dend*_ was computed by equation 4.5.47 of Johnston and Wu:

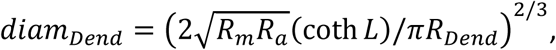

where a fixed value of 173 Ω·cm (Destexhe et al., 1998) for the axial resistivity *R*_*a*_, the dendrite length*L*_*Dend*_ was computed by equation 4.4.15 of Johnston and Wu:

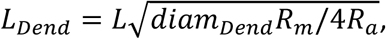

and the passive conductance *g*_*pas*_ was estimated by

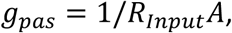

where *A* = *Π*(*diam*_*soma*_^2^ + *diam*_*Dend*_*L*_*Dend*_) is the total surface area of the neuron.

Initial values for *active parameters* were identical for all model neurons and were taken from literature values (Amarillo et al., 2014; Destexhe, 1998) as follows: 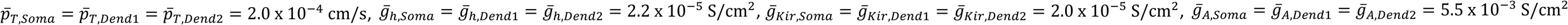 and 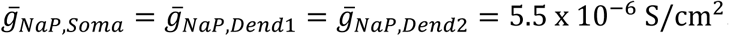.

### Parameter optimization

The voltage responses of a model neuron to a set of 12 different GABA_B_ IPSC waveforms (4 pharmacological conditions at 3 different conductance amplitude scales) was compared with the responses of the corresponding recorded neuron. To emphasize the LTS response, model neurons excluded fast sodium and potassium channels and the experimental responses were median-filtered with a time window of 30 ms to remove action potentials. As multiple objective functions were of interest (Druckmann et al., 2007), a total error *E*_*totat*_ was computed by a weighted average of component errors (Figure 3E):

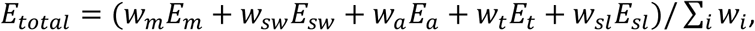

where *E*_*m*_ is the average error across all traces for whether the presence or absence of LTS matches (if an LTS is missed an error of 18 is assigned; if an LTS is falsely produced an error of 6 is assigned), *E*_*sw*_ is the average root-mean-square error of voltage values, *E*_*a*_ is the average LTS peak value error across all LTS-matching traces, *E*_*t*_ is the average LTS latency error across all LTS-matching traces, *E*_*st*_ is the average LTS maximum slope error across all LTS-matching traces, and the *w*_*i*_’s denote the corresponding weights. Based on the maximum LTS latency observed in recorded neurons, the fitting region was restricted to between 0-1.8 seconds after the start of the IPSC. Traces were also allowed to have different sweep weights, in which case the average across traces were computed using root-mean-square.

A modified version of the Nelder-Mead simplex algorithm (Lagarias et al., 1998) was used to update parameter values within defined parameter bounds, with the objective of minimizing *E*_*totat*_. For each simplex, the maximum number of iterations was 2000, the maximum number of error evaluations was 4000, the relative error tolerance was 0.1, the relative parameter change tolerance was 0.1, the coefficient of reflection was *ρ* = 1, the coefficient of expansion was *Χ* = 2, the coefficient of contraction was *γ* = 0.75, the coefficient of shrinkage was σ = 0.8. The initial simplex was set up so that each vertex deviates in a parameter to be varied by *δ* = 2/3 of the parameter’s total range. To transform values to an unconstrained parameter space so that the Nelder-Mead method could be applied, each bounded parameter value was first linearly transformed to the region [–1, 1] than transformed to (–∞, ∞) using the inverse tangent function.

A total of 21 different iterations were run to optimize each model neuron, with the weights among different error types and weights among different traces varying from iteration to iteration. For some poorly fit neurons, the initial parameters for that neuron were replaced by the best-fit neuron’s parameters from the previous iteration. One trace per GABA_B_ IPSC waveform was used during model optimization, but all traces were used for the final model evaluation (Figure 3E).

### Network Models

Each 2-cell network includes one single-compartment model reticular thalamic neuron, described in previous work (Klein et al., 2018), and one of the well-fitted, 3-compartment model thalamocortical neurons described earlier. The reticular thalamic neuron has an AMPA receptor that is synaptically activated by the thalamocortical neuron, and the thalamocortical neuron has a GABA_B_ receptor in the somatic compartment that is synaptically activated by the reticular thalamic neuron. Synaptic currents were evoked with 100% probability and a delay of 1 ms whenever the presynaptic neuron reached a voltage threshold of -30 mV.

AMPA receptors were adapted from Sohal et al. (2000). Reflecting more recent physiological measurements (Deleuze and Huguenard, 2006), AMPA currents were adjusted to bring rise time to 0.5 ms, decay time to 5.6 ms and maximal conductance of 7 nS per synapse. The reversal potential for AMPA currents was maintained at 0 mV.

GABA_B_ receptors were as described before for single model TC neurons. In order for overlapping IPSCs to be summed linearly, we expanded Eq1 into 18 terms, each having its own rise and decay exponential time constants. To be consistent with dynamic clamp experiments, a reversal potential of -115 mV was used in simulations, although the networks behave similarly if a reversal potential of -100 mV was used instead (data not shown). Relative to the values used in dynamic clamp, the conductance amplitudes were scaled by 1/12 to account for temporal summation from an RT burst, and the synaptically-evoked IPSCs was verified to be comparable to recorded values (Figure 7B).

Each 200-cell network assumed a bilayer architecture, with one 100-cell circular layer of single-compartment model reticular thalamic neurons and one 100-cell circular layer of 3-compartment model TC neurons. The identity of the TC neuron was chosen from each of the 31 well-fitted model TC neurons. Each reticular thalamic neuron inhibited the nine nearest TC cells, whereas each TC neuron excited the five nearest reticular thalamic neurons (Sohal and Huguenard, 2003). The passive leak conductance of each model reticular thalamic neuron was randomly selected from a uniform distribution between 45 and 55 µS/cm^2^ (Sohal and Huguenard, 2003). The passive leak conductance of each model TC neuron was randomly selected from a uniform distribution between -10% to 10% of the optimized value.

Network simulations were performed with leak reversal potentials and initial membrane potentials set to a value between -73 mV and -60 mV, at 1 mV increments, so that a total of 14 repetitions were applied for each model network. After a delay of 3 seconds to allow for state variable stabilization, either the reticular thalamic neuron in the 2-cell network or each of the center 20 reticular thalamic neurons in the 200-cell network was injected with a square current pulse (0.2 nA, 40 ms). Simulations were performed with a 0.1 ms integration time step and continued for a total of 30 seconds. Pooling all action potential spikes after stimulation end, we detected bursts and computed an oscillation period and an oscillatory index using the same algorithm as that for multiunit recordings, except for a bin width of 100 ms for spike histograms and a minimum peak prominence of 0.5 relative to the largest secondary peak for the filtered autocorrelation function. Oscillation probability was defined as the proportion of simulations (with varying holding potentials) that induced oscillations with at least three bursts. Percent of active TC cells was defined as the percentage of model TC neurons in the network that produced at least one spike. Half activation latency was defined as the time it took for half of the final percentage of active cells to be activated. Mean oscillation period, mean oscillatory index and mean half activation latency were computed by restricting to trials with a successfully evoked oscillation.

### Statistics

MATLAB R2019b was used for all statistical analysis. Since the number of available data points for LTS and burst features was significantly different for the Dual-Block condition, we performed a paired t-test or a signed-rank test between the Control condition and the Dual Block condition. We used either repeated-measures ANOVA or the Friedman’s test for comparison across the Control, GAT1-Block and GAT3-Block conditions.

Paired comparisons were applied if not otherwise specified. Normality of the differences to the within-subject mean was assess for all groups using a combination of the Lilliefors test, the Anderson-Darling test and the Jarque-Bera test. Normality was satisfied when the geometric mean of three p values was at least 0.05 for all groups. When normality was satisfied, the paired-sample *t*-test was used when there are two groups and repeated-measures ANOVA (with multiple comparison) was used when there are more than two groups. When normality was not satisfied, the Wilcoxon signed-rank test was used when there are two groups and Friedman’s test (with multiple comparison) was used when there are more than two groups. Tests were two-tailed with a significance level of 0.05. Error bars reflect 95% confidence intervals. All violin plots used a bandwidth that was 10% of the maximum data range.

### Drugs

Bicuculline methiodide was purchased from Sigma-Aldrich (St. Louis, MO). The GAT1 blocker NO-711 [(1,2,5,6-tetrahydro-1-[2-[[(diphenylmethylene)amino]oxy]ethyl]-3-pyridinecarboxylic acid hydrochloride] and GAT3 blocker SNAP-5114 [1-[2-[tris(4-methoxyphenyl)methoxy]ethyl]-(S)-3-piperidinecarboxylic acid] were purchased from Tocris Bioscience (Minneapolis, MN).

### Code

Data analysis, statistical tests, NEURON model setup and NEURON model optimization were performed using custom MATLAB code, available online at https://blabuva.github.io/Adams_Functions/. The NEURON .hoc and .mod files used are available on ModelDB.

## Movie Legends

**Movie 1**. Voltage and T-type calcium channel open probability discrepancy trajectories in response to *s*GAT3-Block and *s*Dual-Block GABA_B_ IPSCs.

Simulated traces for model neuron 1 of Figure 3, resampled at 1 ms intervals, in response to either *s*GAT3-Block (red) or *s*Dual-Block (purple), as shown in Figure 5. Traces over simulation time are shown for (**A**) somatic voltage, (**B**) distal dendritic T channel activation gate, (**C**) distal dendritic T channel inactivation gate and (**D**) distal dendritic T channel open probability discrepancy. (**E**) Phase plot of somatic voltage versus distal dendritic T channel open probability discrepancy. (**F**) Phase plot of the concavity versus slope of the distal dendritic T channel open probability discrepancy. All traces are aligned in movie time to the decision points (circles), defined as the point at which the open probability discrepancy curve reaches zero concavity. The time intervals shown in (E-F) correspond to time regions with larger marker sizes in (A-D). Note that the traces for instantaneous and steady-state activation gates in (B) overlap on this scale. Following either *s*GAT3-Block or *s*Dual-Block, the open probability discrepancy curve reaches an inflection point with zero concavity. However, only following the sGAT3-Block does the open probability discrepancy curve reach a significantly positive concavity, drive the open probability discrepancy above threshold (10^−2^) and produce a robust low-threshold spike (LTS). The slope of the open probability discrepancy curve is higher for an LTS-producing trajectory than for an LTS-lacking trajectory.

**Movie 2**. Voltage and T-type calcium channel open probability discrepancy trajectories in response to all GABA_B_ IPSCs.

Same as Movie 1 but in response to all *s*IPSCs shown in Figure 5. (**A-F**) See descriptions for Movie 1. Note that following *s*Control-Block, the maximum concavity for the open probability discrepancy curve is negative, so its trajectory in the open probability discrepancy concavity versus slope phase plot (F) never reaches above the y = 0 line.

**Movie 3**. Voltage and T-type calcium channel open probability discrepancy trajectories in response to GABA_B_ IPSCs with two different decays.

Simulated traces for model neuron 1 of Figure 3, resampled at 1 ms intervals, in response to GABA_B_ IPSCs using the *s*Dual-Block waveform (conductance amplitude scaled by 200%), but with time constants set to 2.0 (orange) or 2.3 (red) seconds. These two time constants correspond to the last LTS success and the first LTS failure, respectively, in Figure 6D. (**A-F**) See descriptions for Movie 1.

**Movie 4**. Voltage and T-type calcium channel open probability discrepancy trajectories in response to GABA_B_ IPSCs with varying time constants.

Same as Movie 3 but in response to all GABA_B_ IPSC time constants shown in Figure 6D. (**A-F**) See descriptions for Movie 1. There appears to be a threshold for the slope of the open probability discrepancy curve that differentiates between an LTS-producing trajectory and an LTS-lacking trajectory.

## Acknowledgements

We thank the University of Virginia Medical Scientist Training Program and the NIH grants R01-NS099586 and R01-NS034774 for funding support. We thank Shin-Shin Nien for help with figure organization. We thank Dr. Peter Klein, Dr. Lise Harbom and Katie Salvati for help with manual LTS scoring. We thank Dr. Farzan Nadim for providing guidance on computational modeling.

